# Image-based spatial transcriptomics identifies molecular niche dysregulation associated with distal lung remodeling in pulmonary fibrosis

**DOI:** 10.1101/2023.12.15.571954

**Authors:** Annika Vannan, Ruqian Lyu, Arianna L. Williams, Nicholas M. Negretti, Evan D. Mee, Joseph Hirsh, Samuel Hirsh, David S. Nichols, Carla L. Calvi, Chase J. Taylor, Vasiliy. V. Polosukhin, Ana PM Serezani, A. Scott McCall, Jason J. Gokey, Heejung Shim, Lorraine B. Ware, Matthew J. Bacchetta, Ciara M. Shaver, Timothy S. Blackwell, Rajat Walia, Jennifer MS Sucre, Jonathan A. Kropski, Davis J McCarthy, Nicholas E. Banovich

## Abstract

The human lung is structurally complex, with a diversity of specialized epithelial, stromal and immune cells playing specific functional roles in anatomically distinct locations, and large-scale changes in the structure and cellular makeup of this distal lung is a hallmark of pulmonary fibrosis (PF) and other progressive chronic lung diseases. Single-cell transcriptomic studies have revealed numerous disease-emergent/enriched cell types/states in PF lungs, but the spatial contexts wherein these cells contribute to disease pathogenesis has remained uncertain. Using sub-cellular resolution image-based spatial transcriptomics, we analyzed the gene expression of more than 1 million cells from 19 unique lungs. Through complementary cell-based and innovative cell-agnostic analyses, we characterized the localization of PF-emergent cell-types, established the cellular and molecular basis of classical PF histopathologic disease features, and identified a diversity of distinct molecularly-defined spatial niches in control and PF lungs. Using machine-learning and trajectory analysis methods to segment and rank airspaces on a gradient from normal to most severely remodeled, we identified a sequence of compositional and molecular changes that associate with progressive distal lung pathology, beginning with alveolar epithelial dysregulation and culminating with changes in macrophage polarization. Together, these results provide a unique, spatially-resolved characterization of the cellular and molecular programs of PF and control lungs, provide new insights into the heterogeneous pathobiology of PF, and establish analytical approaches which should be broadly applicable to other imaging-based spatial transcriptomic studies.

## Main

The human lung is structurally complex, with a diversity of specialized epithelial, stromal and immune cells playing specific functional roles in anatomically distinct locations. Large-scale changes in the structure and cellular makeup of this distal lung is a hallmark of pulmonary fibrosis (PF) and other progressive chronic lung diseases. PF is a progressive syndrome in which functional lung alveoli are lost and replaced by pathological expansion of the extracellular matrix^1^, and can occur in the setting of known environmental or occupational exposures (i.e., asbestosis, silicosis, mold), systemic disorders (connective tissue/autoimmune diseases), monogenic syndromes (short telomere syndromes, surfactant protein mutations, Hermansky-Pudlak Syndrome), or without a clearly established cause (the idiopathic interstitial lung diseases, ILDs, including idiopathic pulmonary fibrosis, IPF)^2^. IPF remains the most common and progressive form of PF; most patients succumb to their disease or require lung transplantation within 3-5 years of diagnosis, and available antifibrotic treatments only modestly slow the inexorable decline of lung function^3,4^.

A hallmark of histopathologic findings in the lungs of patients with IPF (described as “usual interstitial pneumonia”, UIP^5^) is spatial heterogeneity, where extensively remodeled regions of the lung can be found immediately adjacent to regions of relatively preserved alveolar architecture. This spatial variability of disease pathology has been hypothesized to represent asynchronous disease evolution in the lung (“temporal heterogeneity”). In addition to the spatial heterogeneity of pathology, IPF lungs are characterized by: “proximalized epithelial metaplasia”, wherein cell types typically found in conducting airways are observed in the distal lung epithelium; development and accumulation of cystic-appearing structures filled with mucus (“honeycomb cysts”); and the emergence of “fibroblastic foci” (subepithelial collections of fibroblasts) which have been speculated to represent the “leading edge” of disease pathology in the lung^6^. Along with genetic evidence linking IPF susceptibility to the lung epithelium^7–10^ and data from a variety of experimental models, these classical histopathologic features support the prevailing conceptual model of IPF pathogenesis^11^ whereby chronic/recurrent injury to the distal lung epithelium results in dysfunctional alveolar repair and culminates in progressive fibrotic remodeling.

Although the cellular complexity and spatial heterogeneity of disease in the lung presents fundamental challenges when using bulk-tissue methods for genomic analysis, single-cell biology approaches are well-suited for such investigations. Large collaborative studies using droplet-based single-cell RNA-sequencing (scRNA-seq) have refined understanding of the cellular makeup of the normal human lung^12–15^, and a number of studies have highlighted the dramatic changes in the cellular makeup and molecular programs in IPF lungs including a variety of disease-emergent and disease-perturbed cell types and states^10,16–22^. Striking findings have included initial descriptions of a diversity of fibroblast subtypes^10,19,22^, accumulation of epithelial cells in the distal lung co-expressing molecular programs characteristic of AT2 cells and distal airway secretory cells^19^, emergence of *KRT17*+/*KRT5-* “aberrant basaloid” cells^19,20^, expansion of *COL15A1*+ endothelial cells^20^, and changes in macrophage phenotypes including prominent emergence of *SPP1*+ macrophages^17,18,23,24^. The spatial heterogeneity of pathology implies that within a given IPF lung, multiple distinct pathologic programs may be simultaneously occurring in distinct spatial regions (niches); it is thus critical to understand the spatial context within which cellular and molecular programs mediate disease pathogenesis. To this end, we utilized image-based spatial transcriptomics with sub-cellular resolution to investigate the evolution of alveolar niche dysregulation in idiopathic and other forms of PF.

### Diverse cellular landscape of the lung

To characterize the cellular and molecular architecture of the human lung across health and disease, we generated a spatial transcriptomics dataset measuring the expression of 343 genes across 28 3-5 mm lung tissue cores from 6 unaffected and 13 PF donors using the 10X Genomics Xenium platform (**Fig. 1, Supplementary** Fig. 1, and **Supplementary Table 1**). To capture the regional heterogeneity in PF lungs, we profiled paired samples from more (MF) and less (LF) fibrotic regions of 9 PF lungs and samples covering a range of pathologic remodeling (“intermixed”, INT) from 4 individuals, including 3 larger biopsies. Of the 13 PF patients, the most frequent diagnosis was IPF (n = 7, 53.8%) (**Supplementary** Fig. 1). The majority of donors self-reported European ancestry (78.9%), and 8 (42.1%) reported current or prior tobacco use.

**Figure 1.**
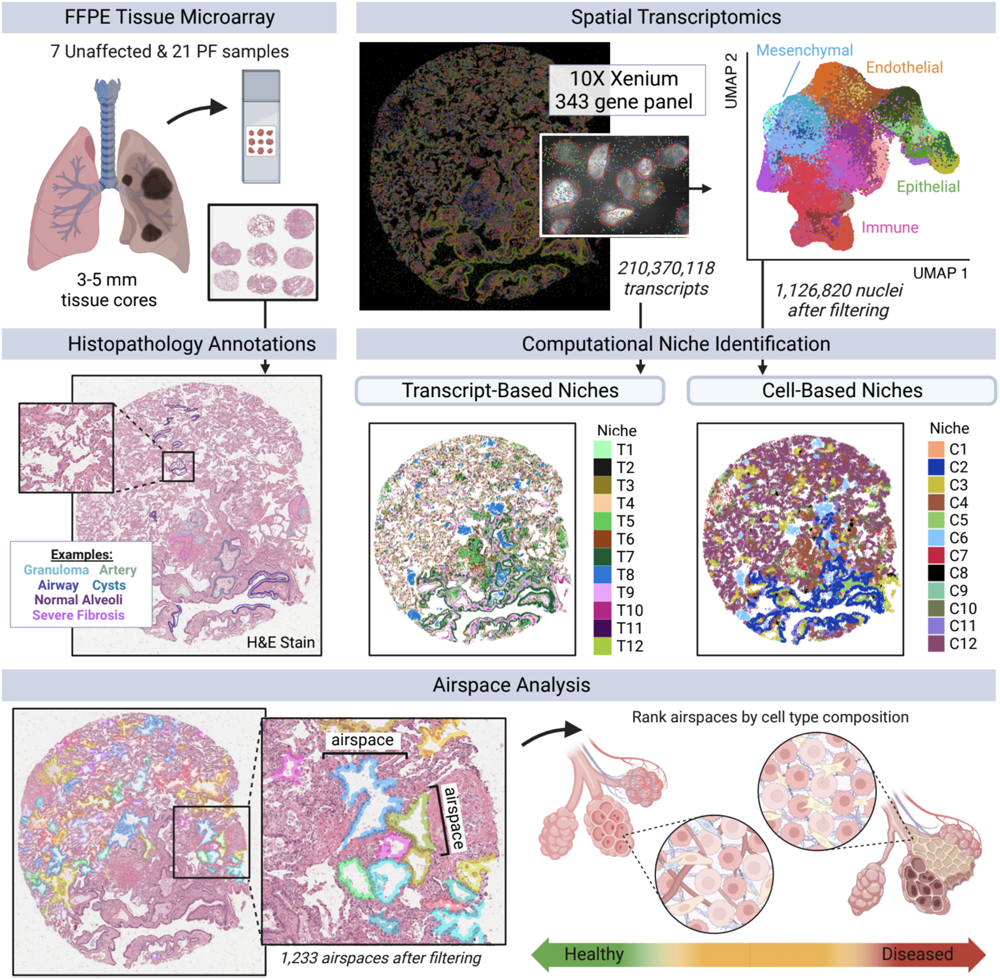
Outline of spatial transcriptomics processing and analysis pipeline. 28 lung tissue cores (3-5 mm) from unaffected and PF donors were processed on tissue microarrays (TMAs) of 3-9 samples each on the Xenium Analyzer instrument. We quantified expression of 343 genes at sub-cellular resolution using a custom panel. After filtering, we retained 210,370,118 high-quality transcripts. After additional filtering, we annotated cell types for 1,126,820 segmented nuclei across the endothelial, epithelial, immune, and mesenchymal lineages.

In total, we measured 210,370,118 high quality transcripts across the 28 samples. To enable spatially-resolved single cell analysis, we partitioned transcripts into cells using automated segmentation to define cell and nuclear boundaries (Fig. 1). As cell segmentation remains a challenge in the field, most notably with a high degree of uncertainty around cellular boundaries and assignment of transcripts to cells, we chose to focus on transcripts overlapping the nuclear boundary. After performing quality filtering (see Methods), we retained 1,126,820 cells that contained 69,406,577 transcripts (Fig. 1). Single-cell processing was carried out using modified versions of standard approaches (see Methods). Overall, we found substantial recovery of a diverse range of cell types, with 39 cell types at our primary annotation level (6 endothelial, 11 epithelial, 13 immune, and 9 mesenchymal; **Supplementary** Fig. 2**, Supplementary** Fig. 3 and **Supplementary Table 2**). Using this annotation approach, we identified recently-described and/or disease-emergent cell types/states including *SCGB3A2+* epithelial cells in terminal airways^25–27^, transitional AT2 cells^19^, *SCGB3A2*+/*SFTPC*+ cells^19,26^, and *KRT5*-/*KRT17*+ “aberrant basaloid” cells^19,20^. We also recovered cell types historically underrepresented in scRNA-seq, including endothelial and mesenchymal cells^10,13^ (**Supplementary** Fig. 2).

Focusing first on establishing the spatial context for cell types/states that have been recently described in scRNA-seq datasets, we observed that consistent with other recent reports^25–27^, *SCGB3A2+* epithelial cells were restricted to small/terminal airways in control lungs, but found widely in remodeled areas of PF lungs where they frequently (but not exclusively) co-expressed *SFTPC*^19^ (**Supplementary** Fig. 4a). As discussed further below, *KRT5*-/*KRT17*+ “aberrant basaloid” cells were absent from structurally normal alveolar regions, and generally found in *SFTPC+/SCGB3A2+* remodeled airspaces. *KRT8*, widely used as a “transitional cell” marker in mouse injury models^28–30^ was expressed relatively broadly in the diseased lung epithelium, but was most mostly highly expressed in *KRT5*-/*KRT17*+ cells (**Supplementary** Fig. 3). Among mesenchymal cells, *PI16+* adventitial fibroblasts were largely associated with vascular structures, while *WNT5A*+ myofibroblasts were observed in ductal regions and at the interface between the lamina propria and adventitia of conducting airways (**Supplementary** Fig. 4b,c). A spectrum of activated fibroblasts expressing varying levels *CTHRC1* and/or *FAP* were most concentrated in subepithelial regions underlying areas of extensive epithelial metaplasia, but were also found more diffusely in some samples. *COL15A1+* endothelial cells were found relatively widely in PF samples with advanced structural remodeling (**Supplementary** Fig. 4d).

We next compared cell type composition across the unaffected, LF, and MF categories using *propeller*^31^, which revealed significant differences in cell type composition across groups for 24 of the 39 identified cell types (61.5%, FDR < 0.05) (**Supplementary** Fig. 5, **6** and **Supplementary Table 3**). Compared to unaffected samples, the disease groups had lower proportions of many cell types typically associated with healthy alveoli, including AT1 and AT2 cells, capillary endothelial cells, alveolar fibroblasts, and *FABP4*+ macrophages. In contrast, LF and MF samples had increased proportions of activated fibroblasts, dendritic cells, B and T-cells, venous endothelial cells, as well as *KRT5-/KRT17+* cells and *MUC5B+* secretory cells. Overall, these results reveal dramatic changes in the cellular composition of the PF lung, corroborating and refining insights from prior studies characterizing the cellular makeup and pathologic diversity of PF lungs^5,16–22,32–37^.

### Cellular and molecular characterization of pathology

The spatial heterogeneity of disease pathology in PF presents the opportunity to directly link gene expression changes to local disease severity as pathology can be simultaneously assessed in the same section from which gene expression was captured. This approach allows us to move beyond categorical designations of disease severity to capture the full spectrum of molecular and cellular dysregulation across patients and samples. Towards this end, we assigned pathology scores for each sample to describe the degree of lung tissue remodeling based on pathology, with higher pathology scores indicative of increased disease severity (Fig. 2a and **Supplementary Table 4**; see Methods). We tested for associations between changes in pathology score and both cell type composition and gene expression using linear regression. We used pseudobulk gene expression values, summing the expression of each gene per sample, including transcripts not segmented into nuclei (see Methods). We found that 19 cell types (48.7%) and 115 genes (33.5%) were significantly associated with pathology score (FDR < 0.01) (Fig. 2b**, Supplementary** Fig. 7, 8, and **Supplementary Tables 5**, **6**). Generally, proportions of capillary and alveolar cells associated negatively with pathology score (**Supplementary** Fig. 7 and **Supplementary Table 5**), as did many of their corresponding marker genes (Fig. 2b, **Supplementary** Fig. 8, and **Supplementary Table 6**). By contrast, higher pathology scores were associated with an increased proportion of activated fibroblasts and all lymphoid cell types. We also performed a cell type-aware analysis, identifying hundreds of genes that were significantly associated with pathology score in individual cell types (**Supplementary** Fig. 9 and **Supplementary Table 7**). Next we sought to characterize the molecular and cellular basis of distinct pathologic features. In addition to generating summative scores reflecting overall disease pathology in a given sample, we annotated representative examples of 21 features across 22 lung samples (Fig. 2a**, Supplementary** Fig. 10, and **Supplementary Table 8**; see Methods), including classical IPF-associated features (i.e. fibroblastic foci, microscopic honeycombing), general pathologic features (muscularized arterioles, granulomas, goblet cell metaplasia, tertiary lymphoid structures), and additional features (i.e. remodeled epithelium, severe fibrosis, mixed inflammation). Characterizing the cellular diversity within these pathologic features then allowed us to establish key molecular processes underlying these disease features (Fig. 2c and **Supplementary** Fig. 11). For example, we observed that regions of granulomatous inflammation contained multiple macrophage subtypes in addition to T-cells (Fig. 2d). Regions of microscopic honeycombing were characterized by a low cuboidal epithelium expressing a variety of airway epithelial cell programs (including multiple secretory cell types as well as basal cells), while regions of goblet cell metaplasia were located in larger airways and notable for expression of *MUC5B* as the predominant airway mucin (Fig. 2d). Tertiary lymphoid structures (TLSs) contained an expected T-cell predominance, while fibroblastic foci included not only activated (*CTHRC1*+) fibroblasts, but multiple fibroblast subtypes, smooth muscle cells, and macrophages (Fig. 2d). Regions of hyperplastic alveolar epithelial remodeling were largely composed of “transitional AT2” cells co-expressing markers of AT1, AT2 and distal airway secretory cells (Fig. 2d).

**Figure 2.**
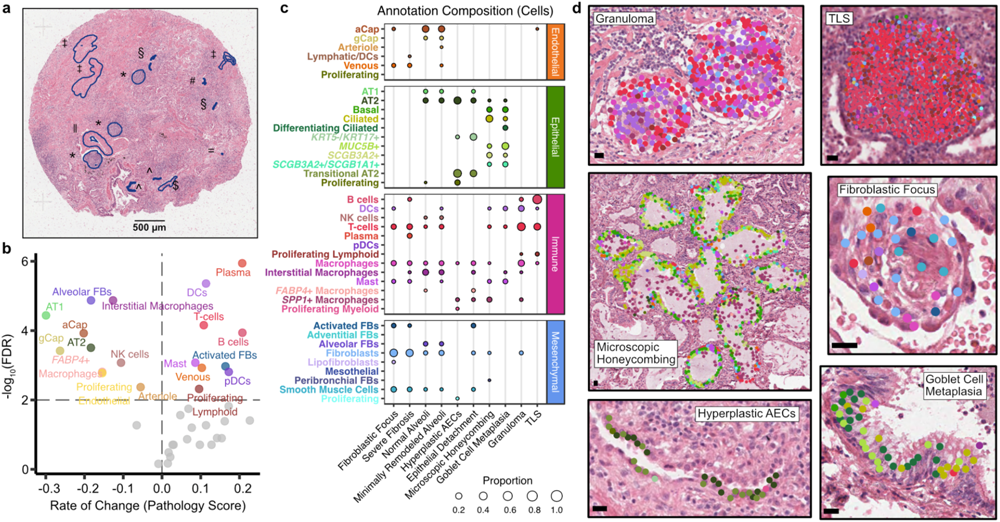
The molecular and cellular basis of clinically relevant PF histopathologies. **a**, Each sample was assigned a pathology score based on a semiquantitative scale assessing the number/severity of pathologic features. Here, we show a representative sample (VUILD107MF; IPF diagnosis) with annotated features. Features are represented by symbols as follows: ǁ muscularized artery; ‡ severe fibrosis; § hyperplastic AECs; # epithelial detachment; = multinucleated cell; ^ remodeled epithelium; $ goblet cell metaplasia; * mixed inflammation. **b**, Cell types significantly associated with pathology score are labeled in the volcano plot. The horizontal and vertical dashed lines show the significance threshold (FDR < 0.01) and split the plot into genes negatively and positively associated with pathology score, respectively. Cell types on the right are present in higher proportions in samples with high pathology scores. **c**, The cell type composition of select annotations of interest, as a proportion of the number of cells across an annotation (each column sums to 1). **d**, Examples of select annotations on H&E images overlain with cell types in the annotated region. Each point represents a cell centroid. Cell type colors are matched to **b**. Scale bars represent 20 µm. Example annotations were taken from the following samples: granuloma - VUILD96MF (sarcoidosis); tertiary lymphoid structure, TLS - VUILD110 (CTD-ILD); microscopic honeycombing - VUILD78MF (IPAF); fibroblastic focus - VUILD105LF (IPF); hyperplastic AECs - VUILD107MF (IPF); goblet cell metaplasia - VUILD104MF (IPF).

### Disease-specific cellular interactions are revealed by niche analyses

We next sought to extend our contextual analyses beyond a-priori defined pathological features with the goal of comprehensively defining spatially-integrated cellular/molecular units in the lung and characterizing their evolution in disease. To this end, we employed two complementary computational approaches to partition samples into regions of molecular and cellular similarity (i.e., spatial “niches”; Fig. 1**, 3a**). First, we used a cell-based approach, building a local neighborhood based on spatial proximity and cell type annotation, followed by k-means clustering as implemented in Seurat v5^38,39^; see Methods). This approach was limited by two factors: 1) it is influenced by choices made as to “granularity” of cell annotation, and 2) it only uses data from transcripts assigned to nuclei. To overcome these limitations, we also developed an approach to identify niches agnostic of cell-assignment by directly using individual transcript data. Using GraphSAGE^38^, we trained a graph neural network model using the spatial location of our transcript data to aggregate local neighborhood information and define an embedding space that provides a new representation for all individual transcripts in our dataset. We then applied Gaussian Mixture Models (GMM) for clustering of transcripts in the embedding space to identify niches. We assigned cells to our transcript-based niches using a consensus approach (see Methods). In both analyses we identified 12 niches (C1-C12 for cell-based clustering and T1-T12 for transcript-based clustering; **Supplementary** Fig. 12), which displayed distinct gene expression signatures (**Supplementary** Fig. 13 and **Supplementary Table 9**) and cell type compositions (Fig. 3b and **Supplementary** Fig. 14). Unaffected samples were primarily defined by healthy alveolar niches (T4, C12) that were substantially depleted in PF samples, and many niches were specific to or enriched in disease, including several immune (T2; C4), fibrotic (T6, T9; C5, C7), transitional epithelial (T3, T7), and airway niches (T1; C2, C11) (Fig. 3c, and **Supplementary** Fig. 12**, 14**). Strikingly, we observed that the niche composition of less fibrotic regions of PF lungs much more closely resembled that of more fibrotic PF than that of control lungs. This unexpected finding implies that despite relative structural preservation, there is extensive molecular pathology in less severely remodeled regions of PF lungs and challenges the paradigm that spatial heterogeneity of pathology reflects a spectrum of the PF biological evolution at the cellular level.

**Figure 3.**
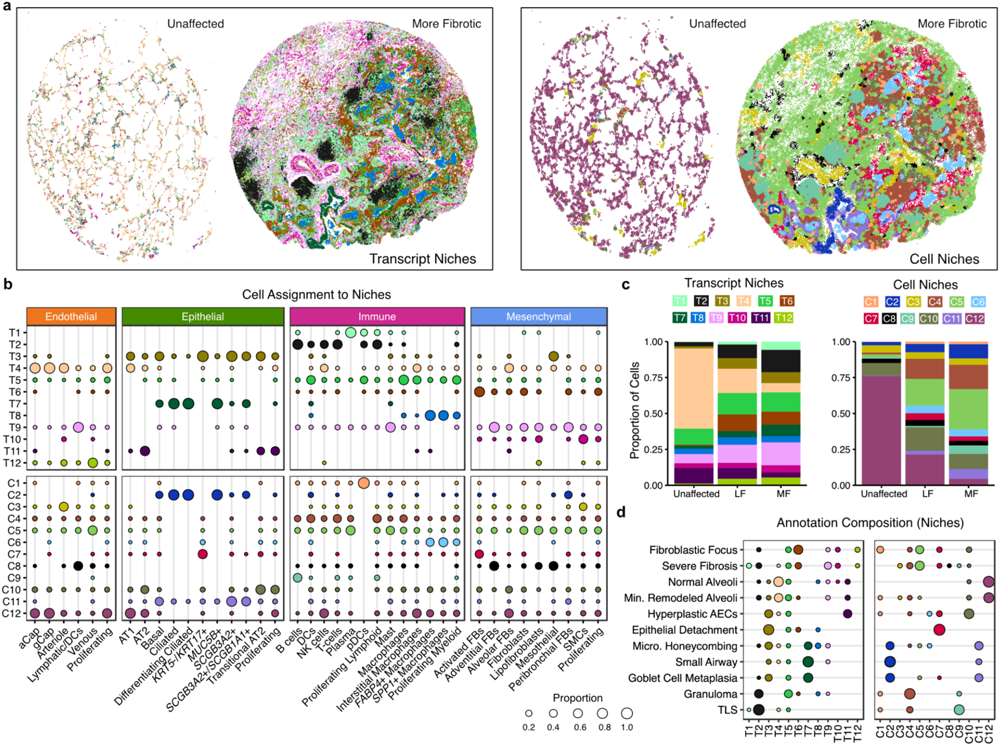
Complementary spatial niche analyses provide comprehensive annotation of tissue remodeling in PF. **a**, Representative examples from both unaffected and PF samples showing transcript-(left) and cell-based niches (right). Shown are VUHD113 and VUILD107MF (IPF diagnosis). For transcript niches, hexbin plots are shown (see Methods). For cell niches, each point is a cell centroid. **b**, Cell assignment to transcript-(top) and cell-based niches (bottom), as a proportion of the number of cells of each type (each column sums to 1). **c**, Bar plots depicting the total proportion of cells across the unaffected, less fibrotic (LF), and more fibrotic (MF) sample types assigned to each transcript and cell niche. **d**, The niche composition of select annotations, as a proportion of the number of cells across an annotation (each row sums to 1). For **b,d,** proportions under 0.01 are not shown.

To directly link the computationally-identified niches to the pathology annotations described above, we characterized the niche composition of each distinct pathology (Fig. 3d and **Supplementary** Fig. 15). Interestingly, some pathologic features were predominantly or near-exclusively found within a single computationally-defined niche. As our pathologic feature annotation was representative but deliberately sparse, this presented the possibility that additional examples of a given pathologic feature would be found within a computationally defined niche. For example, an unexpected feature we observed was patchy epithelial detachment from its underlying basement membrane, which has also been observed in IPF and other forms of PF^40,41^. This feature was predominantly composed of a single niche in both the transcript-and cell-based analysis (T3 and C7 respectively). Surprisingly, these niches contain the vast majority of the fibrosis-associated *KRT5-/KRT17+* (aberrant basaloid) cells identified previously by our group and others^19,20^ (Fig. 3b**,d****, 4a**). Directed by our niche analysis, we identified other regions exhibiting epithelial detachment in additional samples (Fig. 4b). These findings highlight the potential of spatial transcriptomic data to identify specific disease-associated pathologic features directly from the molecular data. Importantly, the niches not only aided in identifying areas of epithelial detachment, but they provided complementary insight into the complex cellular and molecular dynamics underlying this process. The T3 transcript niche predominantly marked a suite of cell types associated with transitional alveolar epithelium including *KRT5*-*/KRT17*+ cells as well as transitional AT2 cells and disease-enriched *SCGB3A2*+ distal airway cells, while the C7 cell niche captured the relationship between *KRT5*-*/KRT17*+ cells and activated fibroblasts expressing *CTHRC1* and *FAP* (Fig. 3b,**d**, **4**).

**Figure 4.**
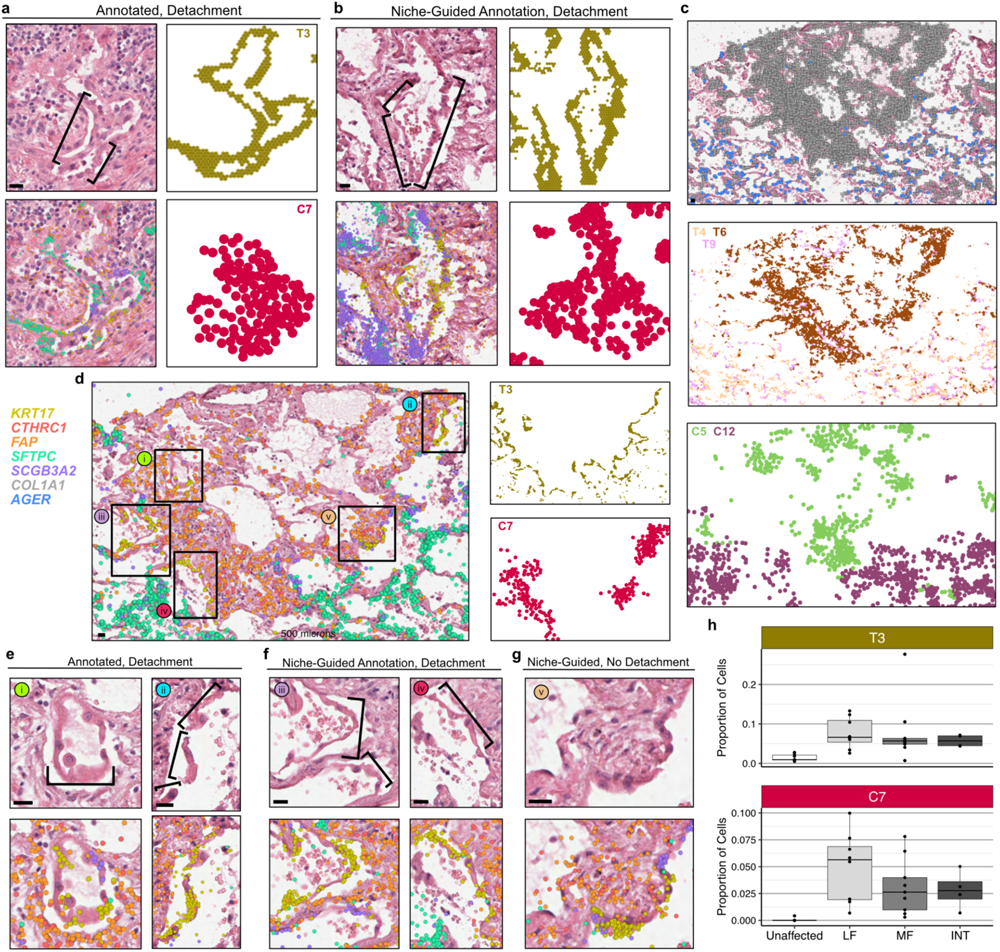
*KRT5*-/*KRT17*+ cells detach at sites of active fibrosis identified by spatial niches. **a**, H&E images of epithelial detachment (denoted by brackets) annotated by a clinician, overlain with transcript expression of the listed genes and compared with the T3 and C7 niches. **b,** an example of epithelial detachment that was not annotated by a clinician but marked by the same niches/genes **c-d,** In one sample, we observed a dense fibrotic region marked by fibrotic niches (T6/T9/C5) and *COL1A1* expression that was lined with epithelial detachment marked by T3 and C7 adjacent to normal alveolar niches (T4/C12) expressing the AT1 cell marker *AGER*. **e**, This region contained two sites of epithelial detachment originally annotated by the clinician (**f**) as well as two additional examples of detaching *KRT5*-/*KRT17*+ and other transitional epithelial cells. **g**, An instance of *KRT5*-/*KRT17*+ cells flanked by activated fibroblasts that were not detaching from the alveolar basement membrane. For **a,b,d-g**, expression is shown for *KRT17*, *CTHRC1*, *FAP*, *SFTPC*, and *SCGB3A2*, for **c**, expression is only shown for *COL1A1* and *AGER*. Scale bars on the bottom left of each H&E represent 20 µm. Samples depicted are as follows: (**a**) VUILD107MF, (**b**) VUILD91MF, and (**c-g**) VUILD91LF, all diagnosed with IPF. **h**, Boxplots showing the proportion of cells assigned to the T3 and C7 niches for each sample, split by sample type - unaffected, less fibrotic (LF), more fibrotic (MF), and intermixed (INT).

In one sample, we observed a striking region of dense fibrosis almost completely lined by the T3 and C7 niches (Fig. 4c,**d**). Adjacent to this area, we found multiple examples of epithelial detachment (Fig. 4e,**f**), but also identified structurally intact epithelium in the same transcriptionally-assigned niche (Fig. 4g). These structurally intact regions appear to be examples of sub-pathologic remodeling, suggesting we can identify molecular and cellular changes that precede histopathology. While the molecular signature of epithelial detachment was prominent at the interface between the putatively advancing fibrotic front (marked by activated fibroblasts) and alveolar epithelium, the larger fibrotic region was marked by stable fibrotic niches and pan-fibroblast marker *COL1A1* (Fig. 4c). Furthermore, we observed open structures reminiscent of alveoli but completely were devoid of epithelium (Fig. 4c**,d**). At a single-point in time, we cannot establish the origin of these “remnant” alveoli, but one possible explanation is they follow epithelial detachment observed at the fibrotic front. Together, these observations raise the possibility that at least some *KRT5*-/*KRT17*+ cells may represent a cell state that precedes epithelial detachment and other progressive pathology. We find the detachment-associated niches are present across disease samples (with slightly increased proportion in LF biopsies) but virtually absent in controls (Fig. 4h).

### Disease-emergent macrophages accumulate in airspaces

In addition to identifying niches which were closely linked to specific pathologic features, our analyses also revealed widespread molecular pathology that did not specifically correspond to classical PF disease features. Indeed, we identified a cell-based niche associated with macrophage accumulation within airspaces that was found across all disease samples irrespective of diagnosis (C6; Fig. 3b**, 5a,b, Supplementary** Fig. 12). While the distinction between remodeled alveolar and distal-airway-derived airspaces is challenging to establish in PF lungs, many of the airspaces marked by these niches appeared to be alveolar in origin, though we observed macrophage accumulation in both alveoli and small airways. Interestingly, the macrophages associated with airspace accumulation appear to include two distinct populations, one marked predominately by *FABP4* and one marked by *SPP1* (Fig. 5c**,d**) which has been described in scRNA-seq datasets as enriched in IPF^18^. Despite their accumulation within airspaces, *FABP4*+ macrophages became less frequent in samples with increasing pathology scores (Fig. 2b and **Supplementary** Fig. 7). Meanwhile, *SPP1*+ macrophages are rarely observed in control lungs, and increased expression of *SPP1* was associated with higher pathology scores (**Supplementary** Fig. 8 and **Supplementary Table 6,7**), suggesting an evolution of macrophage phenotypes characterizes progressive PF. In numerous subjects, discrete regions of *FABP4*+ and *SPP1*+ macrophage accumulation were observed within the same 3-5 mm biopsy. It is not yet clear whether these distinct macrophage subtypes directly promote local remodeling (for example, via *SPP1*-mediated promotion of TGFβ activity^42^), or result from differential polarization related to microenvironmental cues. Indeed, while a number of studies have described phenotypic and compositional changes of macrophages in PF^17,18,23,24^, to our knowledge this is the first integrated, single-cell resolution, spatially-contextualized analysis of macrophage diversity in PF lungs.

**Figure 5.**
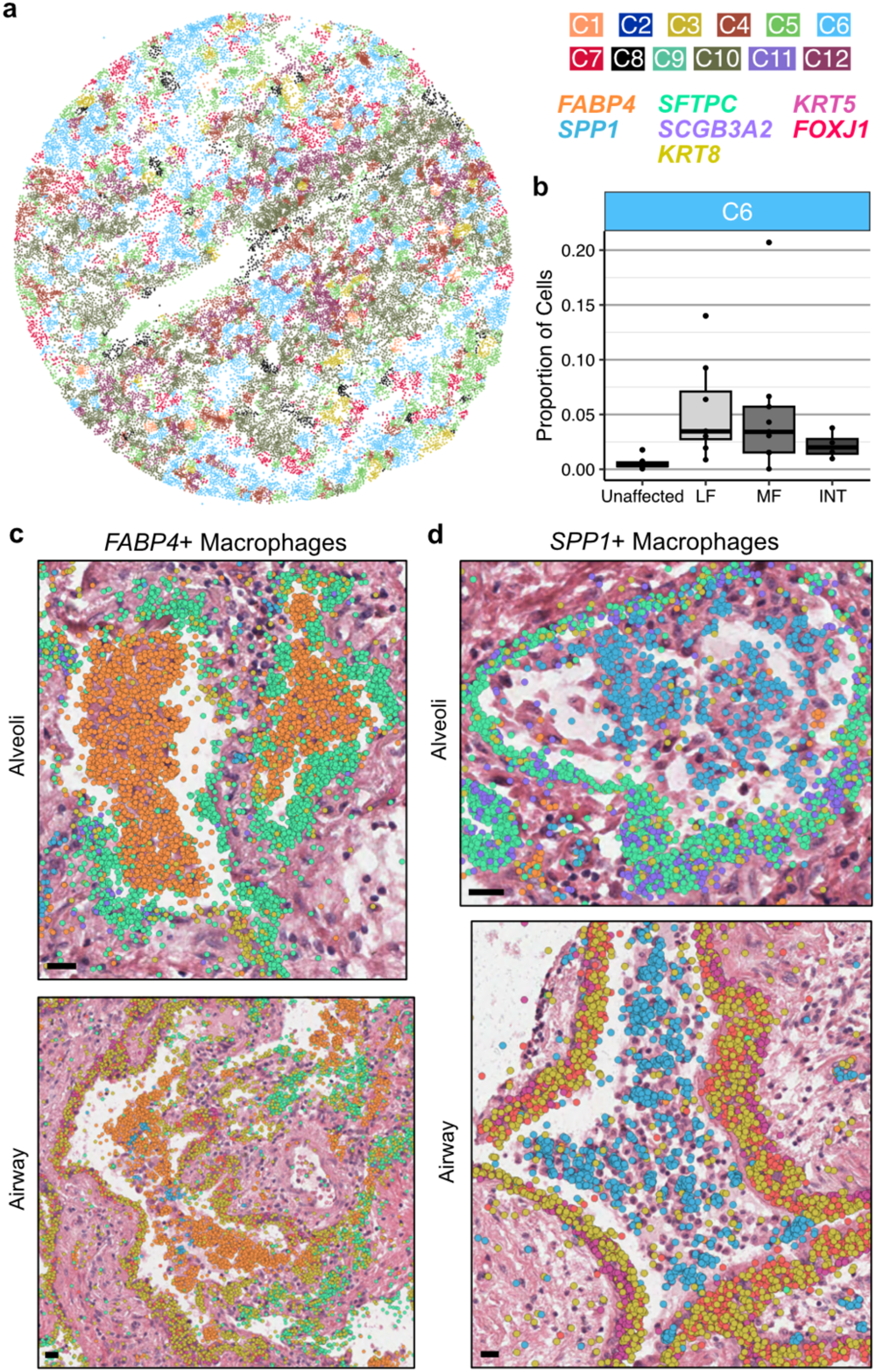
*FABP4*+ and *SPP1*+ macrophages accumulate in PF airspaces and are characterized by a spatial niche. **a**, Representative example of cell niches, including the C6 macrophage accumulation niche (light blue) in sample VUILD102MF (IPF diagnosis). **b**, Boxplot showing the proportion of cells assigned to the C6 niche across sample types, including unaffected, less fibrotic (LF), more fibrotic (MF), and intermixed (INT) samples. **c**,**d**, H&E images of *FABP4*+ (**c**) and *SPP1*+ (**d**) macrophage accumulations in alveoli (top) and airway (bottom), overlaid with transcript expression for listed genes **c** depicts examples from VUILD91MF (top; IPF) and VUILD96LF (bottom; sarcoidosis), and **d** shows samples VUILD78MF (top; IPAF) and VUILD96MF (bottom; sarcoidosis). For both **c** and **d**, all listed genes are potentially visible in each example image if expressed, except *SCGB3A2* which is not shown on the two airway figures. Scale bars on the bottom left of each H&E represent 20 µm.

### A timeline of alveolar dysregulation

The lung’s developmental origin as a branching organ results in a 3-dimensional architecture during which there are ∼22-26 generations of recursively branching conducting airways, followed by terminal bronchioles, respiratory bronchioles, alveolar ducts, and acinar alveoli. In healthy lungs, the abundance of open, alveolar airspaces contributes to the lung’s sponge-like structure. In PF, extensive architectural remodeling and cellular metaplasia result in a diversity of airspaces that approximate the size of an alveolus, but lack a recognizable alveolar epithelium or associated capillary network. As outlined above, even among distal airspaces (i.e. alveoli) of similar size, we observed substantial variation in the degree of cellular and molecular remodeling between the healthy compared to PF lungs, including considerable compositional differences even within different airspaces in a single tissue sample. This led us to hypothesize that across these samples, leveraging the ability to specifically analyze each airspace as an independent unit, it would be possible to capture much of the molecular evolution of alveolar remodeling and begin to establish an “order of events” on the path to end-stage PF.

To this end, we first developed a machine learning approach to identify and segment lumens across samples based on spatial patterns of transcript expression (Fig. 1 and Fig. 6a; see Methods). We then assigned cells to each lumen (see Methods) and filtered the analysis space to include only lumens likely to be alveolar in origin (**Supplementary Table 10**). Next, we ordered the remaining 1,233 airspaces on a continuum of most normal in composition (i.e. “homeostatic”) to most remodeled based on cell type composition using principal component analysis (PCA) and a pseudotime trajectory constructed with *slingshot*^43^ (Fig. 6b). Supporting the validity of this pseudotime strategy, PCA plots show that alveoli from unaffected samples are enriched at the start of the trajectory, and pathology score tends to increase across the trajectory (**Supplementary** Fig. 16). After ordering alveoli by disease severity in this manner, we identified gene expression, cell type composition, and niche proportions significantly associated with pseudotime using generalized additive models (Fig. 6c, **Supplementary** Fig. 16-18 and **Supplementary Tables 11-14**; see Methods). We ordered molecular and cellular changes by the feature’s peak time using the averaged magnitude in sliding windows (**Supplementary Table 15;** see Methods). Supported by both the cell type and niche analyses, we find that initial loss of alveolar homeostasis was marked by a loss of capillary endothelial cells, AT1 cells, alveolar fibroblasts, and interstitial macrophages. The proportion of AT2 cells initially increases (consistent with classical description of “hyperplastic AECs”), but then becomes less frequent as the remaining epithelium has increasing “transitional” and airway-type features/niches. Emergence of activated (*CTHRC1+*) fibroblasts appear around the transition to a more airway-like epithelium, while progressive accumulation of *FABP4+* and then *SPP1+* macrophages are later events (Fig. 6c and **Supplementary** Fig. 16).

**Figure 6.**
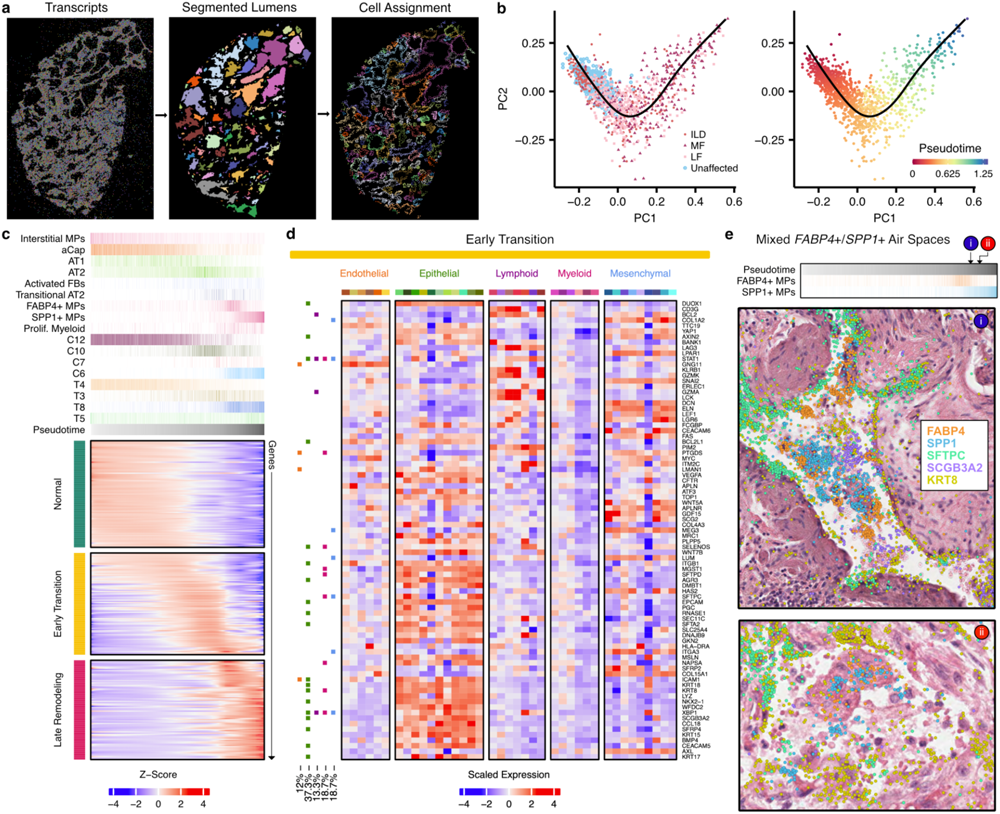
Alveolar remodeling at airspace resolution. **a**, Representation of lumen segmentation pipeline. **b**, Principal components analysis (PCA) and pseudotime analysis projected into the first two PCs, cells are pseudocolored by sample type and pseudotime. **c**, Heatmap of predicted expression of each gene that was significantly associated with pseudotime. The top annotation shows select cell types, cell niches, and transcript niches that were associated with pseudotime, with the darkest shade of each color representing the maximum proportion of that cell type or niche found across all airspaces. **d**, Scaled expression across cell types for the 87 genes with altered expression in the early transition stages in alveolar remodeling from **c**, utilizing only cells that were contained within one of the 1,233 airspaces. Cell type colors match those in Fig. 2a. On the left, boxes are filled in for each gene if it showed a significant change in expression in at least one cell type across the pseudotime of each of the following lineages: endothelial (orange), epithelial (green), lymphoid (purple), myeloid (pink), and mesenchymal (blue). **e,** H&E images of mixed *FABP4*+ and *SPP1*+ macrophage accumulations in two alveoli ranked near the end of the pseudotime trajectory, overlaid with transcript expression for all listed genes. Above the H&Es, each alveolus is marked by its position in pseudotime, and the proportion of *FABP4*+ and *SPP1*+ macrophages is shown for each airspace across pseudotime as in **c**. The example alveoli shown are VUILD115_L1184 (cHP diagnosis; top) and VUILD102_L463 (IPF; bottom). Scale bars on the bottom left of each H&E represent 20 µm.

Focusing on the gene expression data, we manually grouped the expression changes into three broad categories: normal, early transition, and late remodeling. Given limited knowledge of early disease mechanisms, we first focused on the group of genes in the early transition stage. Notably, many of these genes are expressed in both normal and dysregulated airspaces, some being expressed across a large portion of the putatively healthy alveoli (Fig. 6c). We utilized a pseudobulk approach, aggregating gene expression levels across each airspace, leading to associations driven by a mix of molecular and cellular composition changes. To deconvolve these dynamics in the early transition genes, we quantified the expression level in each cell type from all cells within the 1,233 airspaces (Fig. 6d). Furthermore, we carried out a cell type level GAM analysis for 24 cell types with sufficient cells, accounting for cell composition (see Methods). In both our analyses we see a clear localization of signal to epithelial cells, with 37% of the significant associations in the cell type level GAM analysis coming from this lineage (Fig. 6d and **Supplementary Table 16**), and 15.8% localized specifically to AT2 cells. This result is in contrast to the associations identified in the late remodeling category where only 20% of the total associations are from epithelial cells and 37% are associated with myeloid cells (**Supplementary** Fig. 19). Interestingly, when focusing on the myeloid cells during late remodeling, we observed two ordered peaks of *FABP4+* macrophages followed by *SPP1+* macrophages. When we examined airspaces at the intersection of these two peaks, we identified a number of airspaces with both *FABP4+* and *SPP1+* macrophages (Fig. 6e), suggesting the possibility that *FABP4+* macrophage accumulation leads to the recruitment of, or differentiation to, *SPP1+* macrophages. Indeed, prior research has suggested both possibilities^18,44^. Together, these results support a conceptual model where initial events in alveolar remodeling are driven by disruption of the alveolar-capillary interface with activation of regeneration-associated programs in the epithelium, followed by a wave of subepithelial fibroblast activation, then myeloid cell recruitment/proliferation as subsequent events.

## Discussion

In this study, building upon prior lung molecular atlas projects^12–14,19,20,45^, we generated an integrated, spatially-contextualized characterization of the cellular diversity of the adult distal lung in health, and how this evolves in chronic fibrotic lung disease. Leveraging this unique dataset, we developed two complementary strategies which enable quantification and interrogation of multicellular “niches”; we anticipate that these approaches will be broadly applicable to other imaging-based spatial transcriptomic studies and datasets. A key finding emerging from these analyses is that regions of PF lungs that retain relative structural integrity nonetheless have considerable molecular dysregulation. We then developed a novel machine-learning based approach to segment the lung into discrete airspaces and adopted pseudotime trajectory analysis approaches to rank individual airspaces by their degree of pathologic remodeling, quantify the compositional and molecular changes, and establish an ordered sequence of cellular and transcriptional changes that underlies alveolar remodeling in PF.

Beyond contextualizing individual cell types, we established the molecular basis of a diversity of classical histopathologic features of PF. We demonstrated that fibroblastic foci are comprised both activated fibroblasts and several other fibroblast subtypes, frequently also with macrophages; the low cuboidal epithelium in regions of microscopic honeycombing contains several transitional and secretory cell types; “hyperplastic” alveolar epithelial cells reflect a spectrum of mixed alveolar and secretory-cell features. Provocatively, we identified numerous regions where *KRT5-/KRT17+* cells appear to have detached en-masse from their basement membrane when overlying areas adjacent to activated fibroblasts. This finding has been demonstrated previously^40,41^, although there remains debate as to whether this occurs *in-vivo* or reflects an *ex-vivo* artifact. While we cannot conclusively exclude the possibility of *ex-vivo* detachment, we suggest it is relatively unlikely the same feature would be identified in multiple samples, in an analogous cellular and spatial niche context, by chance if this were a stochastic *ex-vivo* artifact. Furthermore, we demonstrate this phenomenon is not driven by sample processing on the Xenium platform by identifying regions of detachment in adjacent sections that did not undergo spatial transcriptomic analysis (**Supplementary** Fig. 20).

The implications of this process are not yet clear, but one possibility is that exposed basement membranes could be prone to patchy fusion, or permit migration of fibroblasts into the airspace where they elaborate pathologic extracellular matrix that leads to eventual airway obstruction, yielding cystic structures distal to the point of fusion/obstruction. In light of recent adoptive transfer studies suggesting IPF basal/basal-like cells can potentiate fibrosis when instilled into the airway^46^, these findings raise the possibility that paracrine effects of *KRT5*-/*KRT17*+ cells could extend beyond immediately adjacent neighbors *in-vivo*.

We also found that whether using cell-aware or cell-agnostic approaches, there appears to be a determinable number of conserved, molecularly-definable spatial “niches” in the human lung. As key cellular processes occur in a spatially and temporally coordinated manner, conceptualizing these niches as distinct functional units allows for directed interrogation of cellular and molecular programs in a specific context. We found that there were substantial shifts in the relative abundance of a given niche across disease pathology. erhaps most strikingly, even within relatively preserved regions of fibrotic lungs, the molecular signature of “normal alveoli” was virtually absent, suggesting that substantial molecular pathology precedes extensive tissue/architectural remodeling.

We then extended this concept further by developing a novel approach to segment individual alveoli/airspaces and explore the evolution of molecular pathology in progressively more remodeled regions. Rather than an initial influx of inflammatory cells or fibroblast activation, these results suggest that disruption of the alveolar epithelium and adjacent capillary network are observed before other structural remodeling is detected. This concept is supported by additional evidence that suggests PF risk is mediated primarily through the lung epithelium^10,47^. Other recognized disease-associated features including emergence of abundant activated fibroblasts and accumulation of macrophages appear to be later events in remodeling. These findings imply that precision therapeutic strategies will likely require concurrent assessment of which cellular mechanisms are most prominent in an individual at a given time; this concept presents potential challenges, but also raises the possibility of improving outcomes (and minimizing toxicities) by better aligning therapeutics with individual patient disease biology.

There are several limitations to this study. First, while this is the largest imaging-based spatial transcriptomic study of the human lung reported to date, this study ultimately reflects a relatively small number of individuals (n=19), samples collected from organ donors or end-stage disease, and samples were obtained from subjects who were predominantly of European ancestry. In the future, we anticipate additional insights will be possible as larger datasets reflecting a broader diversity of subjects and disease are developed. Imaging-based spatial transcriptomic platforms also are inherently semi-targeted, and the probe-set used for this study was informed by prior experience with scRNA-sequencing datasets and developed specifically for cell-identification and examination of a number of established PF-biology related molecular programs and pathways; these platforms are not designed for “unbiased” discovery-based efforts, and it is likely that as these technologies mature, additional refinement of cellular and niche identification will be possible. In addition, cell segmentation remains a challenge, particularly in organs (including the lung) where many cell types have irregular shapes and/or sizes that approach the limits of the optical resolution of the platform. We attempted to mitigate these issues for cell-aware analyses by restricting our dataset to transcripts overlying nuclei, but some degree of “contamination” of transcripts from adjacent/overlying cells remained (**Supplementary** Fig. 3); to this end, differential expression across samples or regions with varying cellular composition (e.g unaffected versus disease) are likely less robust than those possible using sc-or snRNA-seq (**Supplementary** Fig. 9). Initial experience here suggests that transcript-based “niche” analyses offer one approach to overcome these inherent challenges to gain insight into integrated multicellular pathologic processes, and a mix of cell type-agnostic and cell type-aware approaches will be necessary to generate robust results and contextual information.

To summarize, this study provides a comprehensive characterization of the cellular diversity and molecular pathology of the adult distal lung in both health and PF. The identification of conserved molecularly-definable spatial niches and their evolution across disease provides new insights into PF pathogenesis, and the development of novel analytical approaches for quantification and interrogation of multicellular niches using spatial transcriptomic approaches serve as valuable resources for the lung biology community.

## Methods

### Subjects and samples

Peripheral PF lung samples were obtained from lungs removed at the time of lung transplant surgery at VUMC or Norton Thoracic Institute as previously described^10,19,48^. Control lung tissue samples were obtained from lungs declined for organ donation. Diagnoses were determined by the local treating clinicians and affiliated multidisciplinary committee, and confirmed by review of explant pathology according to the current American Thoracic Society/European Respiratory Society consensus guidelines^6,49^. The studies were approved under local institutional review boards (VUMC IRB# 060165,171657 and WIRB# 20181836).

Sample names were generated based on their collection site (Vanderbilt University: “VU”; Translational Genomics Research Institute: “T”), disease status (healthy donor: “HD”; interstitial lung disease “ILD”), unique number assigned to the patient, and where applicable, whether the sample was taken from a less fibrotic (“LF”) or more fibrotic (“MF”) tissue section or a replicate from the same healthy donor (“A” and “B”), e.g. VUHD116A, VUHD113, and TILD117MF.

#### Tissue Microarray (TMA) construction

To maximize efficiency per run, multiple samples could be placed on a single Xenium slide (10.45mmx22.45mm) using a tissue microarray (TMA) design. A 5um section of each lung FFPE block was H&E stained and presented to a physician who identified areas of interest and labeled them as less or more fibrotic (LF or MF) relative to the sample. TMA blocks were designed in a 3×3 or 2×2 pattern for 3mm and 5mm cores, respectively (**Supplementary Table 1**). Sample cores were punched and placed manually using a “Quick-Ray Manual Tissue Microarrayer Full Set” and “Quick-Ray Molds”, according to the manufacturer’s directions.

Empty core spaces were filled with core punches taken from blank paraffin blocks (VWR 76548-194). Once complete, blocks are placed face down on a clean glass slide and briefly heated in a warm drawer (∼45°C) to slightly melt the paraffin together and even the block face. TMA blocks were then cooled to room temperature, removed from the slide, sealed, and stored at 4°C. During TMA preparation, two samples, VUILD105LF and VUHD069, were placed closely together such that a subset of transcripts and nuclei could not be distinguished as belonging specifically to either sample. These transcripts and nuclei were removed from downstream analyses.

### Xenium *in situ* workflow

#### Gene panel design

Xenium in Situ technology requires the use of a pre-defined gene panel. Each probe contains two paired sequences complementary to the targeted mRNA as well as a gene-specific barcode. Upon binding of the paired ends, ligation occurs. The now circular probe is amplified via rolling circle amplification, increasing the signal to noise ratio for target detection and decoding. A total of 343 unique genes were included in the analysis of this dataset. 246 genes came from an early version of the Xenium human lung base panel (PD_336) and 97 genes from a custom designed panel (CVEVZD). The custom panel was curated based on human lung single cell analysis data^19^, selecting for genes useful in cell type identification and/or with suspected involvement in IPF.

#### Xenium sample preparation

As Xenium in Situ technology examines RNA, all protocol workstations and equipment were cleaned using RNase Away (RPI 147002) followed by 70% Isopropanol. All reagents, including water, were molecular grade nuclease-free. Sample preparation began with rehydrating and sectioning FFPE blocks on a microtome (Leica RM2135). 5um sections were placed onto Xenium slides (10X Genomics). Following overnight drying, slides with placed samples were stored in a sealed desiccator at room temperature for ≤10 days. Slides were then placed in imaging cassettes for the remainder of the preparation. Tissue deparaffinization and decrosslinking steps made sub-cellular RNA targets accessible. Gene panel probe hybridization occurred overnight for 18 hours at 50°C (Bio-Rad DNA Engine Tetrad 2). Subsequent washes the next day removed unbound probes. Ligase was added to circularize the paired ends of bound probes (2 hours at 37°C) and followed by enzymatic rolling circle amplification (2 hours at 30°C). Slides were washed in TE buffer before background fluorescence was chemically quenched; autofluorescence is a known issue in lung tissue as well as a byproduct of formalin-fixation ^50^. Following PBS and PBS-T washes, DAPI was used to stain sample nuclei. Finalized slides were stored in PBS-T in the dark at 4°C for ≤5 days until being loaded onto the Xenium Analyzer instrument. Stepwise Xenium FFPE preparation guidelines and buffer recipes can be found in 10X Genomics’ Demonstrated Protocols CG000578 and CG000580.

#### Xenium Analyzer instrument

The Xenium Analyzer is a fully automated instrument for decoding sub-cellular localization of RNA targets. The user marks regions for analysis by manually selecting sample location on an initial low-resolution, full-slide image. After loading consumable reagents and a maximum of two slides per run, internal sample and liquid handling mechanics control experiment progression. Data collection occurs in cycles of fluorescently labeled probe binding, image acquisition, and probe stripping. Images of the fluorescent probes are taken in 4240 x 2960 pixel FOVs. Localized points of fluorescence intensity detected during the rounds of imaging are then defined as potential RNA puncta. Each gene on the panel has a unique fluorescence pattern across the image channels. Puncta that match a specific pattern are then decoded and labeled according to the gene ID. Finally, all image FOVs and associated detected transcripts are computationally stitched together via the DAPI-stained image. Onboard analysis pipelines present quality values for each detected transcript, based on variable confidence in the signal and decoding process. The data in this paper was acquired on instrument software version 1.1.2.4 and analysis version xenium-1.1.0.2. Detailed instructions on instrument operation and consumable preparation can be found in 10X Genomics’ Demonstrated Protocol CG000582.

#### Post-run histology

After the run, slides were removed from the Xenium Analyzer instrument and had quencher removed according to 10X Genomics’ Demonstrated Protocol CG000613. Immediately following, the slides were H&E stained according to the following protocol: xylene (x3, 3min ea), 100% alcohol (x2, 2min ea), 95% alcohol (x2, 2min ea), 70% alcohol (2min), DI water rinse (1min), hematoxylin (1min) (Biocare Medical CATHE), DI water rinse (1min), bluing solution (1min) (Biocare Medical HTBLU-M), DI water rinse (3min), 95% alcohol (30sec), eosin (5sec) (Biocare Medical HTE-GL), 95% alcohol (10sec), 100% alcohol (x2, 10sec ea), xylene (x2, 10sec ea). Coverslipping was performed using Micromount (Leica 3801731) and cured overnight at room temperature. Histology images taken on a 20X Leica Biosystems Aperio CS2.

### Data pre-processing

#### Cell segmentation

Cell segmentation was performed with 10X Xenium onboard cell segmentation (10X Genomics). DAPI stained nuclei from the DAPI morphology image were segmented and boundaries were consolidated to form non-overlapping objects. To approximate cell segmentation, nuclear boundaries were expanded by 15 µm or until they reached another cell boundary.

#### Image registration

Registration was performed between Xenium DAPI morphology images (i.e., nuclei) and H&E stained images of the same slice with the BigWarp plugin implemented in ImageJ (2.14.0)^51,52^. To place both images in the coordinate space of the DAPI morphology image, the H&E stained image was specified as the moving image and the DAPI morphology was specified as the target. Anchors were placed on identifiable landmarks on both images (approx. 200 landmarks per image pair). A thin plate spline warp was then applied to align the corresponding anchor points between the two images. Registered images were manually reviewed for potential visual artifacts resulting from the registration process.

#### Quality filtering and data pre-processing

For each sample, Xenium generated an output file of transcript information including x and y coordinates, corresponding gene, assigned cell and/or nucleus, and quality score. Low-quality transcripts (qv < 20) and transcripts corresponding to blank probes were removed. Seurat v4 was used to perform further quality filtering and visualization on nuclei gene expression data^53^. Nucleus by gene count matrices were created for each sample based on expression of transcripts that fell within segmented nuclei. A single merged Seurat object was created for all samples based on these count matrices and metadata files with nuclei coordinates and area. Nuclei were retained according to the following criteria: ≥ 12 transcripts corresponding to ≥ 10 unique genes, percentage of high-quality transcripts corresponding to blank probes ≤ 5, and nucleus area ≥ 6 and ≤ 80 µm. Because Xenium outputs coordinates based on each slide, which results in samples with shared coordinates across multiple slides, the nuclei coordinates were manually adjusted for visualization so that no samples overlapped during plotting. These adjusted nucleus coordinates were added to the Seurat object as a Dimension Reduction object. Count matrices for transcripts that fell within segmented cells were added to the Seurat object after filtering based on nuclei, though these data were not used for analysis.

#### Dimensionality reduction, clustering, and cell type annotation

Seurat v4 was used to perform dimensionality reduction, clustering, and visualization^53^. Gene expression was normalized per cell using a log1p transformation with the NormalizeData function. Dimensionality reduction was performed with PCA. We selected the number of principal components (PCs) to use by determining the number of PCs that explained more than 90% cumulative variance in the data or the last PC at which the change in explained variance was more than 0.05%, whichever was lower. Nuclei were clustered based on this dimensionality reduction using the Louvain algorithm^54^. Uniform Manifold Approximation and Projection (UMAP) plots of the data were generated using the same number of PCs as used for PCA.

For cell type annotation, we did not rely solely on marker genes to annotate cell types, as some level of gene “contamination” is expected with spatial data because transcripts located in one cell may be assigned to an adjacent cell in sufficiently close proximity. Instead, we incorporated spatial information including cell morphology and histological features to label clusters. Initial Louvain clusters were primarily segregated by cell lineage based on marker genes assessed using the Seurat v4’s *FindMarkers* function, including *PECAM1* (endothelial), *EPCAM* (epithelial), *PTPRC* (immune), *DCN*, *LUM*, and *COL1A1* (all mesenchymal)^53^. New Seurat objects were created for each of the 4 lineages and dimensionality reduction and clustering were performed again as described above. Clusters were then given first-pass cell type labels based on marker genes. The epithelial and immune lineages were further split into subgroups for alveolar and airway cells (epithelial) as well as myeloid and lymphoid cells (immune), and new Seurat objects were generated and re-processed for these subgroups. “Stray” clusters that did not belong to the lineage or subgroup they were originally assigned to were removed and then re-analyzed with the correct grouping. New Seurat objects were then created for each lineage and subgroup based on the first-pass annotations and re-processed. As lineage and subgroup splitting became more accurate, clusters were re-labeled with revised annotations. Final annotations for stray clusters were revised based on shared marker genes with confidently-labeled clusters. Annotation was performed at 6 levels of detail, with 59 cell types at the finest level. Similar cell types were consolidated into broader labels using a bottom-up approach to create the other annotation levels. Subsequent analyses utilized second-broadest annotation level, which consisted of 39 cell types, including 6 endothelial, 11 epithelial, 13 immune, and 9 mesenchymal (**Supplementary** Fig. 2, **3** and **Supplementary Table 2**). For one cell cluster, markers of two cell types (dendritic immune cells and lymphatic endothelial cells) were present and had high spatial overlap. These were labeled “Lymphatic/DCs” and placed in the endothelial lineage.

### H&E image annotation

#### Pathology score assignment

Registered H&E images were viewed in QuPath version 0.4.3^55^ and assessed for the presence of 21 pathologic features, including 4 epithelial (Alveolar: emphysema, hyperplastic alveolar epithelium; Airway: bronchiolar metaplasia, microscopic honeycombing), 1 vascular (muscularized artery), 3 immune (granuloma, tertiary lymphoid structure (TLS), mixed inflammation), and 2 mesenchymal (fibroblastic focus, severe fibrosis) (**Supplementary Table 4**, **8**). For each feature, a score from 0 to 3 (absent, mild, moderate, severe) was assigned to each sample, and the score for all features on a given sample was summed to constitute a pathology score. For samples from 5mm tissue cores, pathology scores were adjusted to account for increased tissue area compared to 3mm samples by multiplying the raw scores by a factor of 0.36.

#### Annotation of histological features

We annotated representative examples of 21 histological features across 22 samples, including 10 epithelial (General: epithelial detachment, remodeled epithelium; Alveolar: hyperplastic alveolar epithelium, interlobular septum, minimally-remodeled alveoli, normal alveoli, remnant alveoli; Airway: goblet cell metaplasia, small airway, microscopic honeycombing), 4 vascular (artery, muscularized artery, vein, venule), 3 immune (granuloma, TLS, mixed inflammation), 3 mesenchymal (airway smooth muscle, fibroblastic focus, severe fibrosis), and 2 general (giant cell, multinucleated cell) feature types (**Supplementary Table 4**, **8**). Annotations were marked on the registered H&E images using QuPath version 0.4.3^55^. Cells were then assigned to annotated regions using a custom Python script. Briefly, annotated regions were scaled by a factor of 0.2125 µm/pixel to harmonize units with the cell centroid coordinates. Cells were then assigned a boolean value for each annotated region corresponding to whether or not the cell centroid fell within the annotated region.

### Comparison to scRNA-seq datasets

We compared cell lineage recovery from the present spatial dataset to two scRNA-seq sources. The first was Natri et al. (2023), a recent sc-eQTL study from our lab^10^, and the other was the Human Lung Cell Atlas (HLCA), which included aggregated data from multiple lung scRNA-seq studies^13^. Studies from the HLCA varied in the disease studied and whether the dataset contained control and/or lung disease samples. We narrowed the scope of our comparison to studies (lung atlases) or samples (sc-eQTL study) that did not specifically enrich or deplete cells from a specific lineage in their pre-processing and did not exclusively contain data from nasal samples. We included 47 samples from the sc-eQTL study (19 control, 28 ILD) and 14 datasets from the HLCA (**Source Data Supplementary** Fig. 2). For each data source, the proportion of cells from each lineage (endothelial, epithelial, immune, and mesenchymal) was calculated first for all samples and then for control and disease samples separately. If an annotated cell type did not have a clear lineage association, it was removed from the analysis. Sample-level data from the present study was compared to sample-level data from the sc-eQTL study and dataset-level information from HLCA.

### Lumen segmentation and airspace identification

#### Initial lumen segmentation

To segment individual lumens from the spatial sequencing data, a custom GPU accelerated image processing algorithm was developed using *scikit-image* (version 0.19.3), RAPIDS cuCIM (version 22.12.00), and CuPy (version 11.2.0) in python (version 3.9.7) to assign unique identifiers to lumens in the spatial transcriptome data^56–58^. Briefly, the location of each detected transcript, excluding transcripts associated with immune cells, were binarized to a 2D image followed by a series of dilation, erosion, and closing operations to define the location of the tissue and segment the lumens. The outer boundaries of the tissue were defined using Alpha Shapes (*alphashape* package version 1.3.1) on a denoised, closed shape representing the entire tissue. Unique identifiers were then assigned to the negative space (lumens) using scikit-image and metrics were calculated for each individual lumen. Cell centers (nuclei) within a cutoff distance, defined as the thickness of a normal alveolar wall, were then assigned to the unique identifier of the closest lumen by searching for the nearest label in a restricted zone using the K-D Tree nearest-neighbor lookup algorithm implemented in *scikit-image*. All cells that were not close to a lumen were given an identifier of zero.

#### Quality filtering to isolate alveolar airspaces

Lumens were first filtered by size to remove false positive segmentations. We retained lumens that contained between 25 and 500 cells and for which the maximum distance between any two nuclei in the lumen was at least 125 µm. To isolate alveoli from endothelial and airway structures, lumens were then grouped k means clustering (k = 7) based on their cell type proportions. Clusters with low proportions of alveolar cell types (AT1, AT2, transitional AT2, and alveolar fibroblasts) and high proportions of non-capillary endothelial (arteriole, venous, and lymphatic) or airway cells (basal, ciliated, differentiating ciliated, *MUC5B*+, and *SCGB3A2*+/*SCGB1A1*+) were removed. To further filter out airway structures, lumens with ≥ 20% of cell assigned to the cell-based airway niches (C2 + C11) were also removed.

#### Ranking alveoli along a pseudotime by severity of tissue remodeling

PCA and pseudotime analysis were performed on the remaining 1,233 alveolar airspaces based on cell type composition to rank them along a continuum of disease severity and tissue remodeling. We converted the cell type counts to proportions and applied PCA on the cell type proportions, and the first three principal components were used for pseudotime analysis using *slingshot*^43^. Briefly, five clusters of alveoli were derived using Gaussian Mixture Model (GMM) clustering, and the minimum spanning tree that connected the cluster centroids was constructed which was then smoothed using simultaneous principal curves. Pseudotime for each alveolus was inferred by projecting the alveolus onto the smooth curve.

#### Assessing changes in cell types, niches, and gene expression along pseudotime

To find cell types, niches, and genes with varied abundances or expression along the pseudotime, we applied generalized additive models using the fitGAM function from *tradeSeq* ^59^ on the aggregated feature counts across lumens under a negative binomial distribution with total cell counts per lumen as an offset. We subsequently applied *associationTest*^59^ for testing the association of features with pseudotime. Cell types or niches were excluded if they were not present in > 10% of segmented alveoli. For heatmap visualizations, the ordering of features (row ordering of the heatmaps) was determined by the pseudotime value of the lumen in which each feature peaked. The peaking lumen for each feature was determined using the sliding window-smoothed values (window size n = 50 for gene expression heatmap, n = 10 for niche and cell type proportion heatmaps). Features with earlier peak times were plotted in the top rows.

##### Classifying gene expression patterns along pseudotime

We clustered fitted gene expression patterns along pseudotime-ordered lumens using GAM models by hierarchical clustering (k = 14) after taking z-scores per gene. We manually separated genes with unimodal and bimodal dynamic expression patterns by visual inspection of the resulting clusters.

##### Altered gene expression along pseudotime within each cell type

We tested which genes had altered expression along pseudotime within a specific cell type using GAMs. We tested 24 cell types with ≥ 3 cells in more than 30 lumens. For each cell type, we aggregated their gene expression across segmented lumens. Within a specific cell type, for each gene’s expression u, we fit a negative binomial GAM model to model the gene count across lumens using pseudotime and cell type proportions as covariates, using the *gam* function with cubic regression splines “bs=’cr’” implemented in mgcv^60^ with

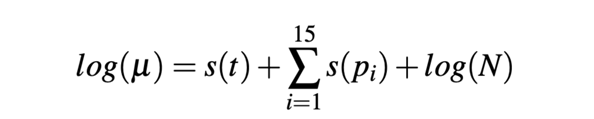

where t represents pseudotime, p_i_ represents proportion of cell type i, and N represents the number of aggregated cells per lumen. We applied the same strategies in knot selection as in *tradeSeq*^59^ and selected 5 knots per smooth term. We included proportions of 15 cell types which had more than 3 cells in more than 100 lumens as smooth terms in addition to the pseudotime term to control for the contamination in detected gene expression signals. Genes that had significant pseudotime terms after multiple testing correction (FDR < 0.01) were summarized and plotted.

### Computational niche identification

#### Transcript-based niches

Transcript-based niches were characterized using a graph neural network model GraphSAGE^38,61^ that directly modeled the detected transcripts of each sample without cell segmentation. GraphSAGE learns the structures in the graph data via sampling and aggregating neighbors, and scales well to large graphs. The list of detected transcripts from each sample was used as input for building the sample graph after removing low-quality transcripts (qv < 20) and blank probes. For each sample, a graph was constructed individually with nodes as individual transcripts and edges drawn between nodes that have euclidean distances (based on spatial x, y locations) smaller than a threshold (*d* = 3.0) similar to a previous study^61^. The same threshold *d* was used for each sample. Each node was also associated with an input feature vector that was the one-hot-encoding of the node’s gene label. Connected components of nodes with fewer than 10 nodes were removed.

We applied a 2-hop GraphSAGE model that for a set of randomly sampled “root nodes” sampled 20 and 10 neighboring nodes at the first-hop and second-hop neighbors respectively to learn an embedding vector of dimension 50 at each hop for each node in the graph. The embedding vector from the last layer of each node integrated information of the node itself and its 2-hop neighbors. To embed all nodes from all samples into the same embedding space, a joined graph consisting of subgraphs from all samples were used as training inputs. To reduce memory usage, we generated a subgraph for each sample from the original sample graph with each subgraph containing 5,000 randomly sampled root nodes along with their 3-hop neighbors as well as the existing edges among them, and these subgraphs were merged into a joint graph (18,594,542 nodes and 421,316,086 edges) for model training. The model parameters were trained in an unsupervised manner by solving a binary classification task for “positive” and “negative” node pairs, in which the model predicts whether or not two nodes should have an edge connection, based on their local neighborhood structures. We trained the model on the merged graph with ten epochs achieving a training accuracy of 0.82^38^.

We obtained the embedding representation of all nodes across 28 samples using the trained 2-hop GraphSAGE model and performed unsupervised clustering on all nodes from all samples (206,449,064 total nodes) using GaussianMixtureModel (k = 12) as implemented in the PyCave library (https://github.com/borchero/pycave).

##### Transcript-based niche plots and assignment of nuclei to niches

We created a hex bin summarization plot for each sample using all transcripts with a *bin_width* 5 for generating the transcript-based niche plots to overcome overplotting issues. Each bin was labeled and colored with the major node label among the transcripts falling in the bin area. Bins with fewer than 10 transcripts were not included in the plots. To assign cells to transcript-based niches, we assigned each cell to its closest hex bin by calculating the euclidean distance of each cell centroid to the hex bin centroids.

#### Cell-based niches

Seurat v5’s^39^ *BuildNicheAssay* function was used to partition cells into 12 spatial niches using k-means clustering based on the cell type composition of their 25 closest neighboring cells.

### Differential expression and composition analyses

#### Differential cell type composition by disease severity

We transformed the cell type proportion per sample using logit transformation and tested whether cell type proportion was different among three disease groups (unaffected, less fibrotic (LF), and more fibrotic (MF) samples.) using the *propeller.anova* function implemented in *propeller*^31^.

#### Representative gene detection in transcript-based niches

We used the linear model framework implemented in *propeller*^31^ and *limma*^62,63^ to test differences in gene proportions across niches and thereby identify representative genes of each niche. We first investigated the proportion of transcripts assigned into each niche within each sample. Samples were excluded from testing for a niche when too few transcripts from that sample (proportion < 5e-4) were assigned into the niche. Gene proportions were logit-transformed, and a linear model was fitted to each gene that modeled the mean transformed gene proportion in each niche while accounting for sample differences. Differentially-abundant genes were then derived by contrasting the mean gene proportion from one niche with the average mean proportion from the other niches.

#### Differential expression, cell type composition, and niche proportions by pathology score

We used the same *propeller*^31^ and *limma*^62,63^ framework for finding gene, cell type, and niche proportions that correlated with changes in pathology scores across samples. Proportions were logit-transformed and linear models were fit to model the relationship between the transformed proportions and pathology scores to find features that varied significantly with pathology scores (FDR < 0.01).

#### Differential expression analysis across annotated pathology features

We aggregated gene counts from cells included in each pathology annotation instance and detected differentially expressed genes among pathology annotations by fitting linear models implemented in *limma*^62,64^. All pathology annotations were included in the analysis except giant cells, because these cells are included in the granuloma annotation. Gene expression was converted to CPM (counts per million) and log_2_ transformed after adding a 0.5 pseudocount. Lowly expressed genes (log_2_CPM < 8) in 50% of the annotation instances were excluded. The *limma voom* function was applied to model the mean-variance relationship and assign weights to each log_2_CPM observation value, which was subsequently used in the linear regression model to account for heteroskedasticity in count data. The number of cells aggregated per annotation instance was added as a covariate. Differentially-expressed genes (DEGs) for each annotation type were then detected through comparing the gene expression per annotation type with the rest. Separately, we also compared epithelial annotations to each other to identify dysregulated genes in pathologic epithelium.

## Data and code accessibility

Raw and processed data are deposited at the Gene Expression Omnibus (GEO) under accession number GSE250346. Custom R, Python, and bash scripts for this project are available on Github at https://github.com/Banovich-Lab/Spatial_PF.

## Acknowledgements

This work was supported by NIH R01HL145372(JAK/NEB), R01HG011886 (NEB/DJM), W81XWH1910415 (NEB/JAK), T32HL094296 (AMS), R01160551 (CMS), P01HL092870 (TSB), Francis Family Foundation (JJG), Vanderbilt Faculty Research Scholars (JJG), K08143051 (JMS), R01HL168556 (JMS), the Department of Veterans Affairs (TSB, JAK), T32HL094296 (ASM), NHMRC GNT1195595 and GNT1162829 to D.J.M, NIH HL158906 and HL126176 (LBW)

## References

1. Hewlett, J. C., Kropski, J. A. & Blackwell, T. S. Idiopathic pulmonary fibrosis: Epithelial-mesenchymal interactions and emerging therapeutic targets. Matrix Biol. 71-72, 112–127 (2018).

2. Lederer, D. J. & Martinez, F. J. Idiopathic Pulmonary Fibrosis. N. Engl. J. Med. 378, 1811–1823 (2018).

3. Richeldi, L. et al. Efficacy and safety of nintedanib in idiopathic pulmonary fibrosis. N. Engl. J. Med. 370, 2071–2082 (2014).

4. King, T. E., Jr et al. A phase 3 trial of pirfenidone in patients with idiopathic pulmonary fibrosis. N. Engl. J. Med. 370, 2083–2092 (2014).

5. Liebow, A. A., Carrington, C. B., Simon, M. & Potchen, E. J. Frontiers of pulmonary radiology. Alveolar diseases: the (1969).

6. Travis, W. D. et al. An official American Thoracic Society/European Respiratory Society statement: Update of the international multidisciplinary classification of the idiopathic interstitial pneumonias. Am. J. Respir. Crit. Care Med. 188, 733–748 (2013).

7. Thomas, A. Q. et al. Heterozygosity for a surfactant protein C gene mutation associated with usual interstitial pneumonitis and cellular nonspecific interstitial pneumonitis in one kindred. Am. J. Respir. Crit. Care Med. 165, 1322–1328 (2002).

8. Seibold, M. A. et al. A common MUC5B promoter polymorphism and pulmonary fibrosis. N. Engl. J. Med. 364, 1503–1512 (2011).

9. Allen, R. J. et al. Genome-wide association study across five cohorts identifies five novel loci associated with idiopathic pulmonary fibrosis. Thorax 77, 829–833 (2022).

10. Natri, H. M. et al. Cell type-specific and disease-associated eQTL in the human lung. bioRxivorg (2023) doi:10.1101/2023.03.17.533161.

11. Pardo, A. & Selman, M. Idiopathic pulmonary fibrosis: new insights in its pathogenesis. Int. J. Biochem. Cell Biol. 34, 1534–1538 (2002).

12. Travaglini, K. J. et al. A molecular cell atlas of the human lung from single-cell RNA sequencing. Nature 587, 619–625 (2020).

13. Sikkema, L. et al. An integrated cell atlas of the lung in health and disease. Nat. Med. 29, 1563–1577 (2023).

14. Guo, M. et al. Guided construction of single cell reference for human and mouse lung. Nat. Commun. 14, 4566 (2023).

15. Schupp, J. C. et al. Integrated Single-Cell Atlas of Endothelial Cells of the Human Lung. Circulation 144, 286–302 (2021).

16. Xu, Y., et al. Single-cell RNA sequencing identifies diverse roles of epithelial cells in idiopathic pulmonary fibrosis. JCI Insight 1, e90558 (2016).

17. Reyfman, P. A. et al. Single-Cell Transcriptomic Analysis of Human Lung Provides Insights into the Pathobiology of Pulmonary Fibrosis. Am. J. Respir. Crit. Care Med. 199, 1517–1536 (2019).

18. Morse, C. et al. Proliferating SPP1/MERTK-expressing macrophages in idiopathic pulmonary fibrosis. Eur. Respir. J. 54, (2019).

19. Habermann, A. C. et al. Single-cell RNA sequencing reveals profibrotic roles of distinct epithelial and mesenchymal lineages in pulmonary fibrosis. Sci Adv 6, eaba1972 (2020).

20. Adams, T. S. et al. Single-cell RNA-seq reveals ectopic and aberrant lung-resident cell populations in idiopathic pulmonary fibrosis. Sci Adv 6, eaba1983 (2020).

21. Heinzelmann, K. et al. Single-cell RNA sequencing identifies G-protein coupled receptor 87 as a basal cell marker expressed in distal honeycomb cysts in idiopathic pulmonary fibrosis. Eur. Respir. J. 59, (2022).

22. Tsukui, T. et al. Collagen-producing lung cell atlas identifies multiple subsets with distinct localization and relevance to fibrosis. Nat. Commun. 11, 1920 (2020).

23. Serezani, A. P. et al. Multi-Platform Single-Cell Analysis Identifies Immune Cell Types Enhanced in Pulmonary Fibrosis. Am. J. Respir. Cell Mol. Biol. (2022) doi:10.1165/rcmb.2021-0418OC.

24. Ayaub, E. A. et al. Single Cell RNA-seq and Mass Cytometry Reveals a Novel and a Targetable Population of Macrophages in Idiopathic Pulmonary Fibrosis. bioRxiv 2021.01.04.425268 (2021) doi:10.1101/2021.01.04.425268.

25. Basil, M. C. et al. Human distal airways contain a multipotent secretory cell that can regenerate alveoli. Nature 604, 120–126 (2022).

26. Kadur Lakshminarasimha Murthy, P., et al. Human distal lung maps and lineage hierarchies reveal a bipotent progenitor. Nature 604, 111–119 (2022).

27. Rustam, S. et al. A Unique Cellular Organization of Human Distal Airways and Its Disarray in Chronic Obstructive Pulmonary Disease. Am. J. Respir. Crit. Care Med. 207, 1171–1182 (2023).

28. Strunz, M. et al. Alveolar regeneration through a Krt8+ transitional stem cell state that persists in human lung fibrosis. Nat. Commun. 11, 3559 (2020).

29. Kobayashi, Y. et al. Persistence of a regeneration-associated, transitional alveolar epithelial cell state in pulmonary fibrosis. Nat. Cell Biol. (2020) doi:10.1038/s41556-020-0542-8.

30. Wang, F. et al. Regulation of epithelial transitional states in murine and human pulmonary fibrosis. J. Clin. Invest. 133, (2023).

31. Phipson, B. et al. propeller: testing for differences in cell type proportions in single cell data. Bioinformatics 38, 4720–4726 (2022).

32. Katzenstein, A. L. Pathogenesis of ‘fibrosis’ in interstitial pneumonia: an electron microscopic study. Hum. Pathol. 16, 1015–1024 (1985).

33. Gracey, D. R., Divertie, M. B. & Brown, A. L., Jr. Alveolar-capillary membrane in idiopathic interstitial pulmonary fibrosis. Electron microscopic study of 14 cases. Am. Rev. Respir. Dis. 98, 16–21 (1968).

34. Blumhagen, R. Z. et al. Spatially distinct molecular patterns of gene expression in idiopathic pulmonary fibrosis. Respir. Res. 24, 287 (2023).

35. Eyres, M. et al. Spatially resolved deconvolution of the fibrotic niche in lung fibrosis. Cell Rep. 40, 111230 (2022).

36. Joshi, N. et al. A spatially restricted fibrotic niche in pulmonary fibrosis is sustained by M-CSF/M-CSFR signalling in monocyte-derived alveolar macrophages. Eur. Respir. J. 55, (2020).

37. Chilosi, M. et al. Abnormal Re-epithelialization and Lung Remodeling in Idiopathic Pulmonary Fibrosis: The Role of ΔN-p63. Lab. Invest. 82, 1335–1345 (2002).

38. Hamilton, W. L., Ying, R. & Leskovec, J. Inductive Representation Learning on Large Graphs. (2017).

39. Hao, Y. et al. Dictionary learning for integrative, multimodal, and scalable single-cell analysis. bioRxiv 2022.02.24.481684 (2022) doi:10.1101/2022.02.24.481684.

40. Zaizen, Y. et al. Alveolar Epithelial Denudation Is a Major Factor in the Pathogenesis of Pleuroparenchymal Fibroelastosis. J. Clin. Med. Res. 10, (2021).

41. Selman, M. & Pardo, A. Role of epithelial cells in idiopathic pulmonary fibrosis: from innocent targets to serial killers. Proc. Am. Thorac. Soc. 3, 364–372 (2006).

42. Kramerova, I. et al. Spp1 (osteopontin) promotes TGFβ processing in fibroblasts of dystrophin-deficient muscles through matrix metalloproteinases. Hum. Mol. Genet. 28, 3431–3442 (2019).

43. Street, K. et al. Slingshot: cell lineage and pseudotime inference for single-cell transcriptomics. BMC Genomics 19, 1–16 (2018).

44. The impact of the lung environment on macrophage development, activation and function: diversity in the face of adversity. Mucosal Immunol. 15, 223–234 (2022).

45. Madissoon, E. et al. A spatially resolved atlas of the human lung characterizes a gland-associated immune niche. Nat. Genet. 55, 66–77 (2023).

46. Jaeger, B. et al. Airway basal cells show a dedifferentiated KRT17highPhenotype and promote fibrosis in idiopathic pulmonary fibrosis. Nat. Commun. 13, 5637 (2022).

47. Liu, Q. et al. The Genetic Landscape of Familial Pulmonary Fibrosis. Am. J. Respir. Crit. Care Med. (2023) doi:10.1164/rccm.202204-0781OC.

48. Bui, L. T. et al. Chronic lung diseases are associated with gene expression programs favoring SARS-CoV-2 entry and severity. Nat. Commun. 12, 4314 (2021).

49. Raghu, G. et al. Diagnosis of Idiopathic Pulmonary Fibrosis. An Official ATS/ERS/JRS/ALAT Clinical Practice Guideline. Am. J. Respir. Crit. Care Med. 198, e44–e68 (2018).

50. Davis, A. S. et al. Characterizing and Diminishing Autofluorescence in Formalin-fixed Paraffin-embedded Human Respiratory Tissue. J. Histochem. Cytochem. 62, 405–423 (2014).

51. Schindelin, J., et al. Fiji: an open-source platform for biological-image analysis. Nat. Methods 9, 676–682 (2012).

52. Bogovic, J. A., Hanslovsky, P., Wong, A. & Saalfeld, S. Robust registration of calcium images by learned contrast synthesis. in 2016 IEEE 13th International Symposium on Biomedical Imaging (ISBI) (IEEE, 2016). doi:10.1109/isbi.2016.7493463.

53. Integrated analysis of multimodal single-cell data. Cell 184, 3573–3587.e29 (2021).

54. Blondel, V. D., Guillaume, J.-L., Lambiotte, R. & Lefebvre, E. Fast unfolding of communities in large networks. J. Stat. Mech. 2008, P10008 (2008).

55. Bankhead, P. et al. QuPath: Open source software for digital pathology image analysis. Sci. Rep. 7, 16878 (2017).

56. van der Walt, S. et al. scikit-image: image processing in Python. PeerJ 2, e453 (2014).

57. RAPIDS Development Team. RAPIDS: Libraries for End to End GPU Data Science. https://rapids.ai/.

58. Okuta R, Unno Y, Nishino D, Hido S, Loomis C. CuPy: A NumPy-Compatible Library for NVIDIA GPU Calculations. in Proceedings of Workshop on Machine Learning Systems (LearningSys).

59. Van den Berge, K., et al. Trajectory-based differential expression analysis for single-cell sequencing data. Nat. Commun. 11, 1–13 (2020).

60. Wood, S. N. Generalized Additive Models: An Introduction with R, Second Edition. (CRC Press, 2017).

61. Partel, G. & Wählby, C. Spage2vec: Unsupervised representation of localized spatial gene expression signatures. FEBS J. 288, 1859–1870 (2021).

62. Ritchie, M. E. et al. limma powers differential expression analyses for RNA-sequencing and microarray studies. Nucleic Acids Res. 43, e47 (2015).

63. Phipson, B., Lee, S., Majewski, I. J., Alexander, W. S. & Smyth, G. K. ROBUST HYPERPARAMETER ESTIMATION PROTECTS AGAINST HYPERVARIABLE GENES AND IMPROVES POWER TO DETECT DIFFERENTIAL EXPRESSION. Ann. Appl. Stat. 10, 946–963 (2016).

64. Law, C. W., Chen, Y., Shi, W. & Smyth, G. K. voom: Precision weights unlock linear model analysis tools for RNA-seq read counts. Genome Biol. 15, R29 (2014).

