## Supplementary Figures for "Image-based spatial transcriptomics identifies molecular niche dysregulation associated with distal lung remodeling in pulmonary fibrosis"

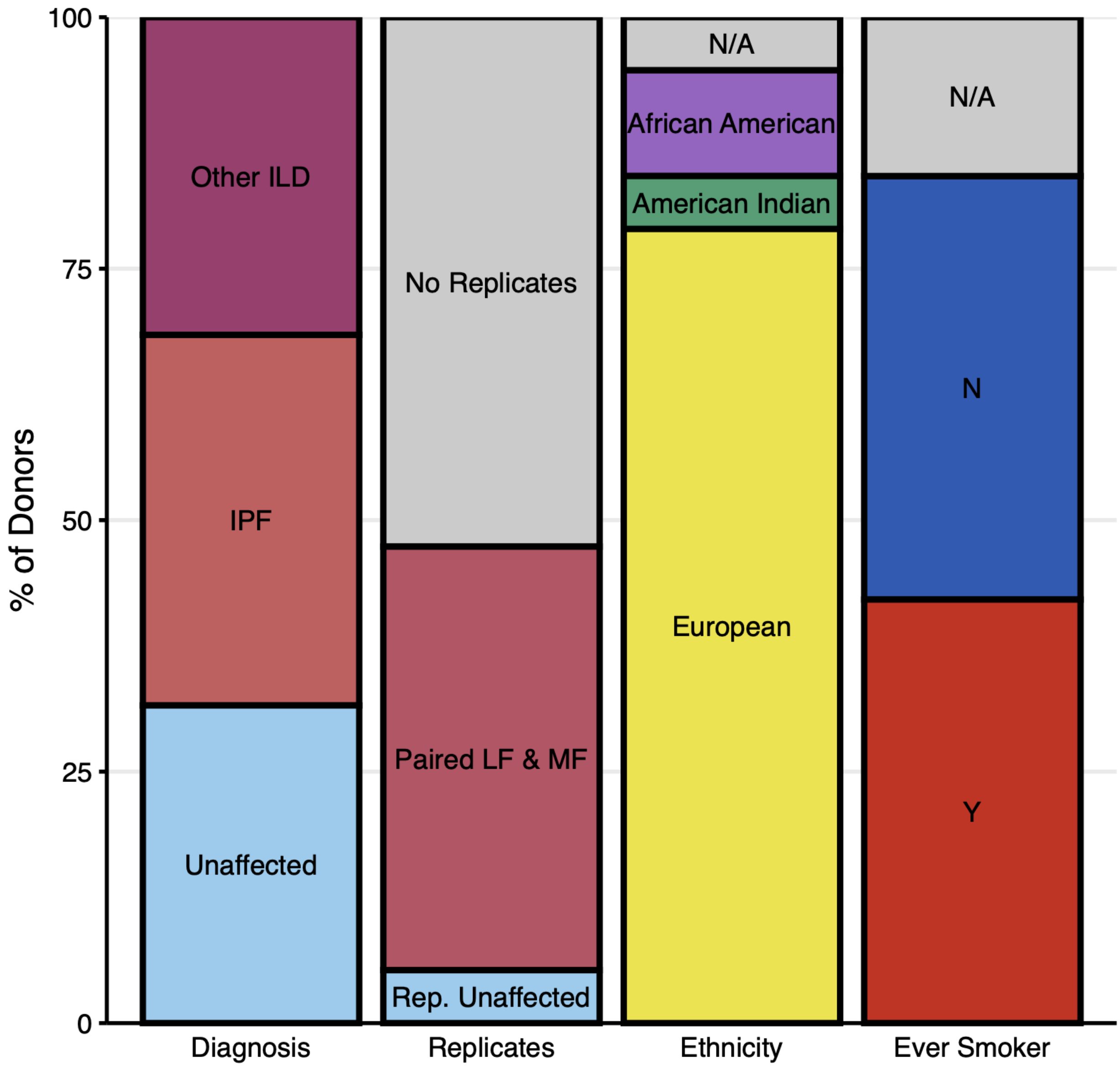


**Supplementary Figure 1: Donor demographic information.**
Samples (N = 28) were collected from 6 unaffected and 13 pulmonary fibrosis (PF) donors across several interstitial lung diseases (ILDs). Most PF donors were diagnosed with idiopathic pulmonary fibrosis (IPF). For some donors, we collected 2 replicate samples for testing. For PF donors with replicate samples, we collected tissue punches from both less fibrotic (LF) and more fibrotic (MF) sections of the same tissue for comparison. Demographic information is also provided for reported ethnicity and smoking status.

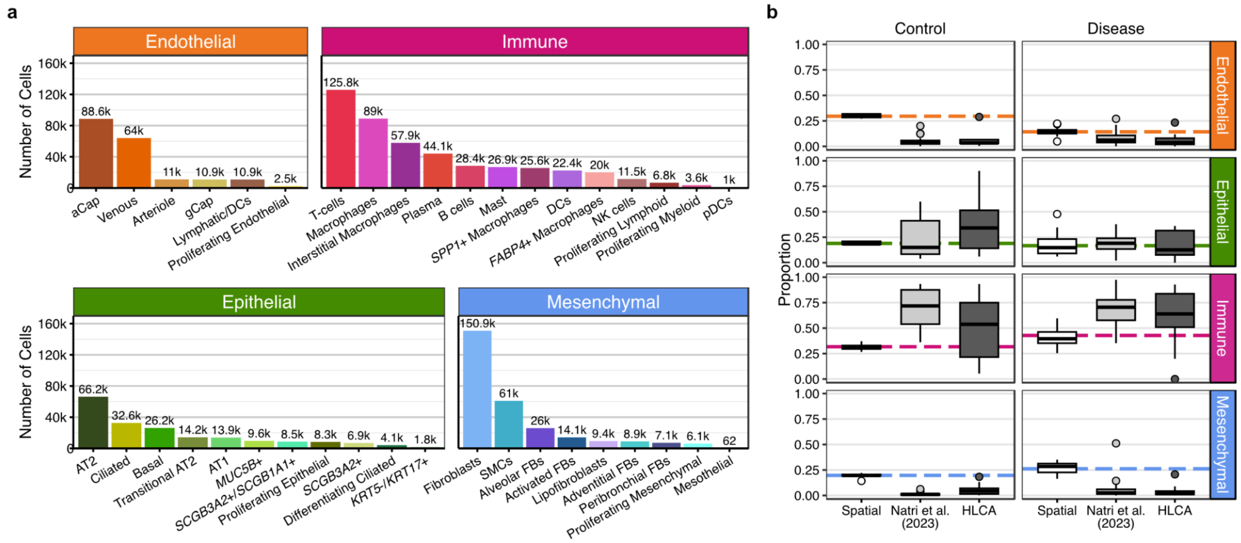


**Supplementary Figure 2: Spatial transcriptomics accurately recovers cell types in PF lung.**

**a**, The frequency of each cell type broken down by all samples. **b**, Comparison of cell type recovery between the present spatial dataset and prior single-cell RNA-sequencing (scRNA-seq) datasets from Natri et al. (2023)[1] and the Human Lung Cell Atlas (HLCA)[2]. Proportion of cells from each lineage was calculated per sample for the spatial dataset and Natri et al. (2023) and per dataset for HLCA. For HLCA calculations, only studies that did not enrich or deplete for any cell lineage were included in the plot. Dashed lines indicate the overall proportion of cells in each lineage for the spatial dataset.

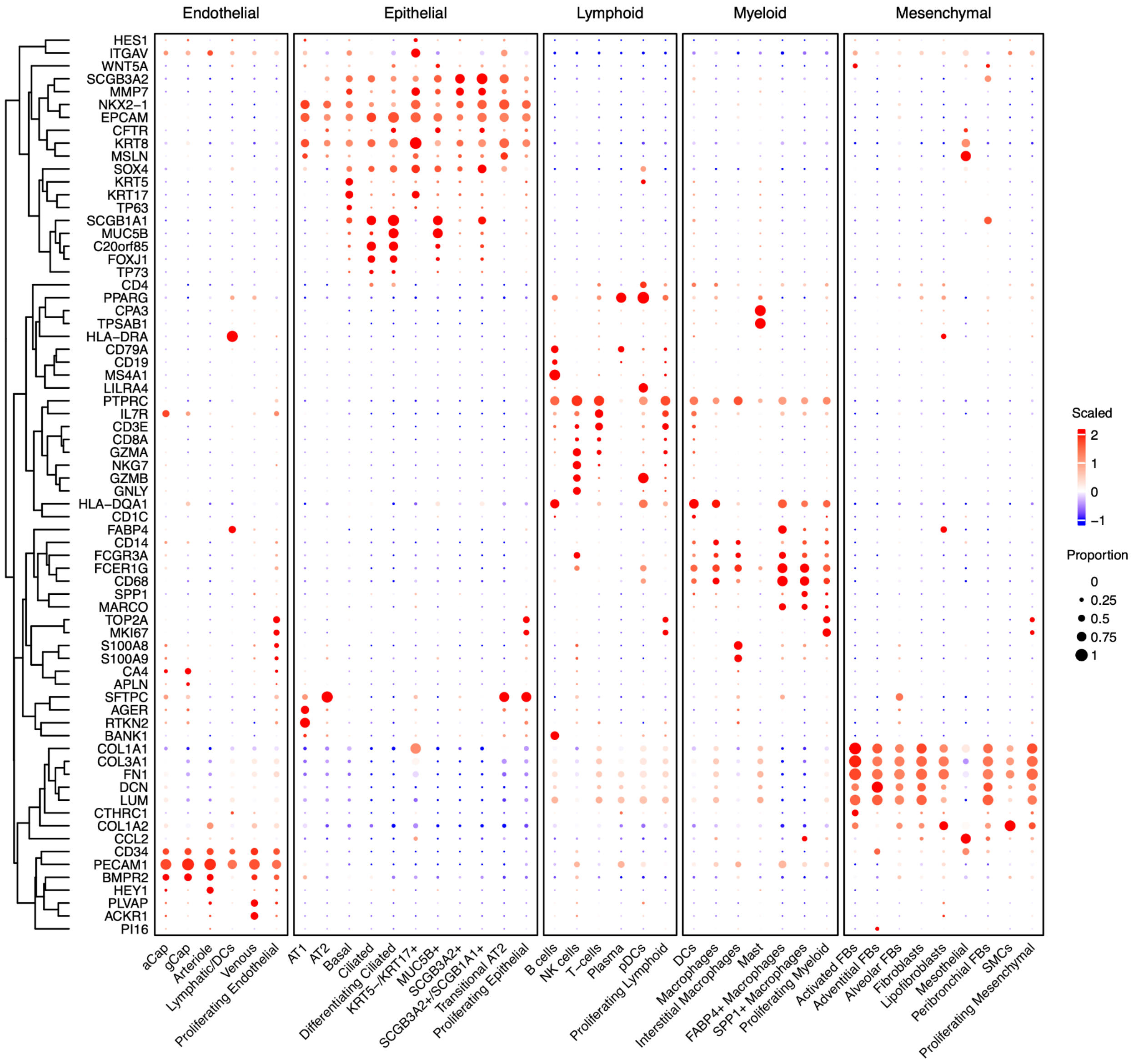


**Supplementary Figure 3: Marker genes for all cell types.**

Of the 343 genes, 70 select genes are shown here to demonstrate cell type annotation of the 39 identified cell types.

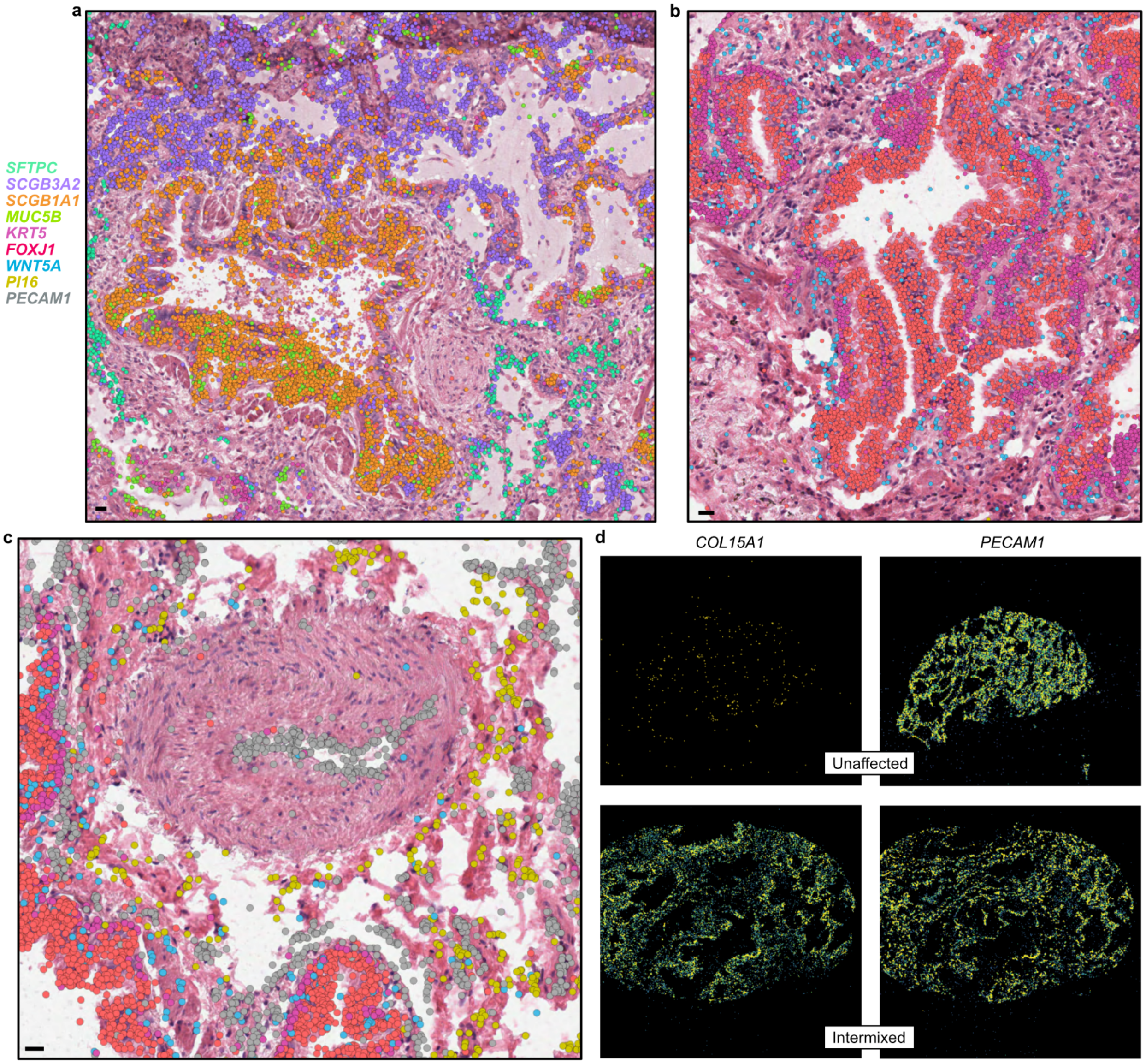


**Supplementary Figure 4: Localization of key cell types.**

**a-c**, Representative examples of (**a**) pathologic *SCGB3A2*+/*SCGB1A1*- positive airways near *SCGB3A2*+/*SCGB1A1*+ airways and *SCGB3A2*+/*SFTPC*+ cells in remodeled alveoli, (**b-c**) *WNT5A*+ (myo)fibroblasts surrounding conducting airways, and (**c**) *PI16*+ adventitial fibroblasts near vasculature. Transcript expression is overlain on H&E images. Legend on top left of figure applies to **a-c**, with the following exceptions: *SFTPC*, *SCGB3A2*, and *MUC5B* are only shown in **a**; *WNT5A* and *PI16* are only shown in **b** and **c**; *PECAM1* is only in **c**. Scale bars represent 20 µm. **d**, Density plots showing expression of disease-enriched endothelial marker *COL15A1* and pan-endothelial marker *PECAM1* in representative unaffected and intermixed (INT) PF samples. Samples depicted are as follows: (**a**) VUILD78MF (IPAF diagnosis), (**b-d**) TILD175 (IPF), and (**d**) VUHD095 (unaffected).

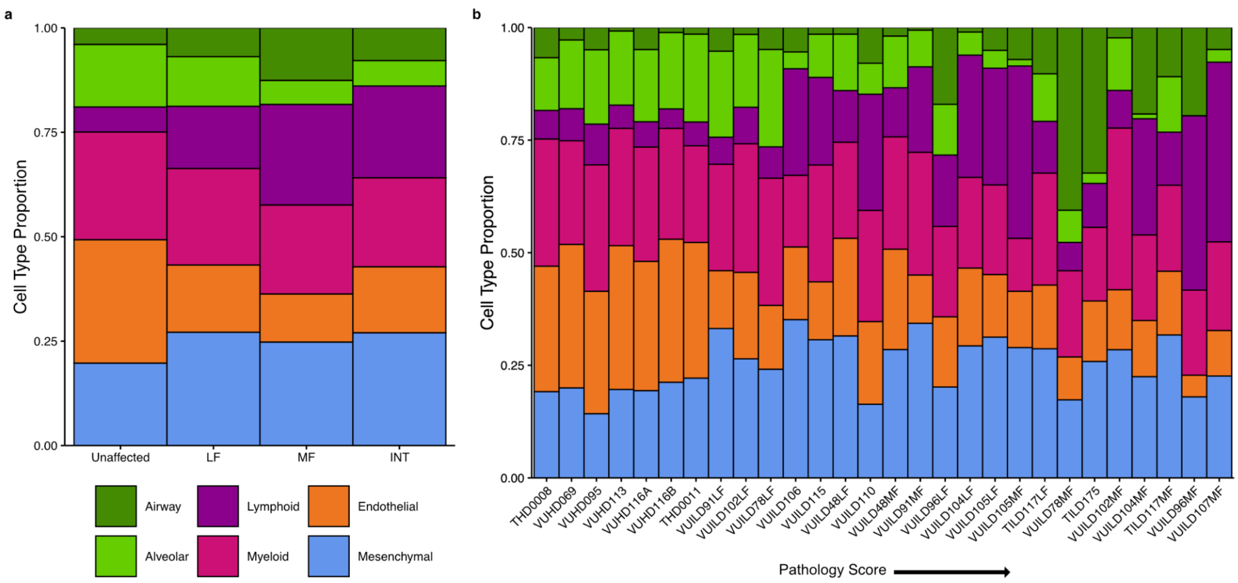


**Supplementary Figure 5: Distribution of cell types across samples.**
Cell type proportions across (**a**) sample type and (**b**) sample, ordered on the x axis by pathology score from lowest to highest. Here, cell types were sorted into broad categories for ease of visualization.

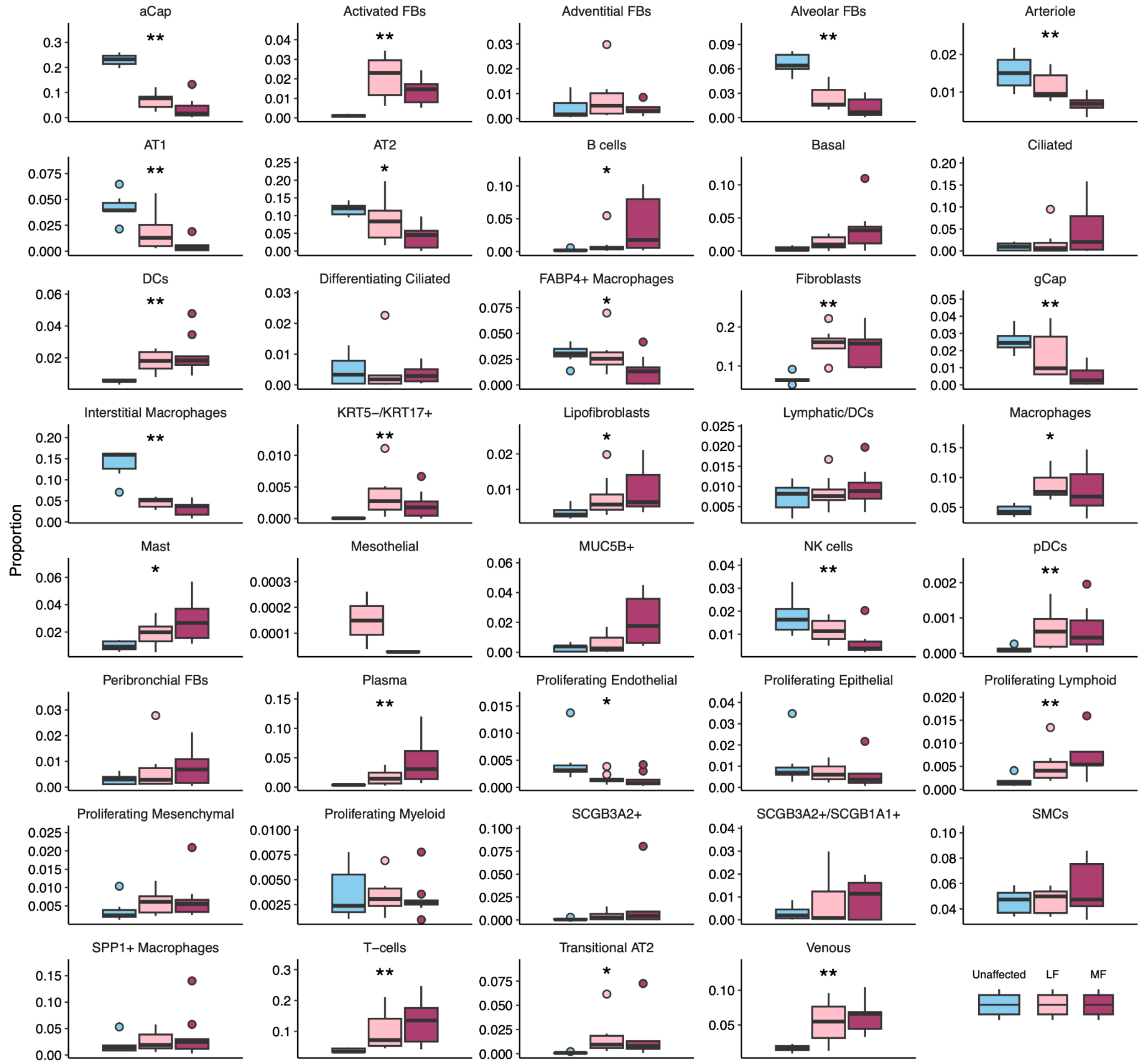


**Supplementary Figure 6: Distribution of cell types by disease severity.**
Boxplots show the proportions of each cell type across unaffected, less fibrotic (LF), and more fibrotic (MF) samples. ** and * indicate significant differences across sample types (FDR < 0.01 and 0.05, respectively), via ANOVA tests as implemented in *propeller* [3].

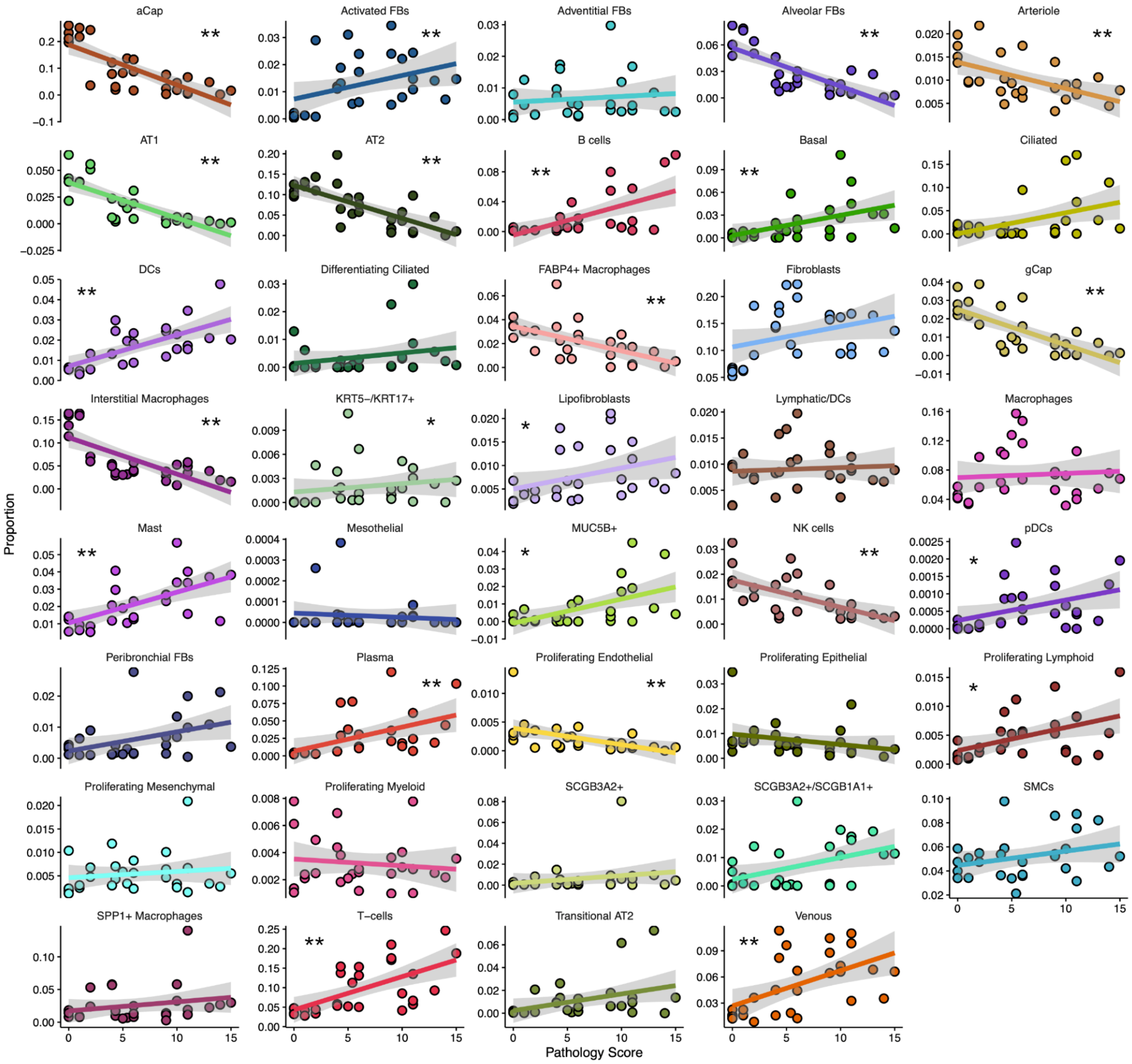


**Supplementary Figure 7: Changes in cell type composition with pathology score.** A linear model framework with *propeller*[3] and *limma*[4,5] was used to assess changes in cell type proportions with changes in pathology score across samples. Proportions were logit-transformed for this analysis; here, correlations of raw proportion with pathology score are shown for simplicity. * and ** represent FDR < 0.01 and < 0.05, respectively, under the linear model (see Methods).

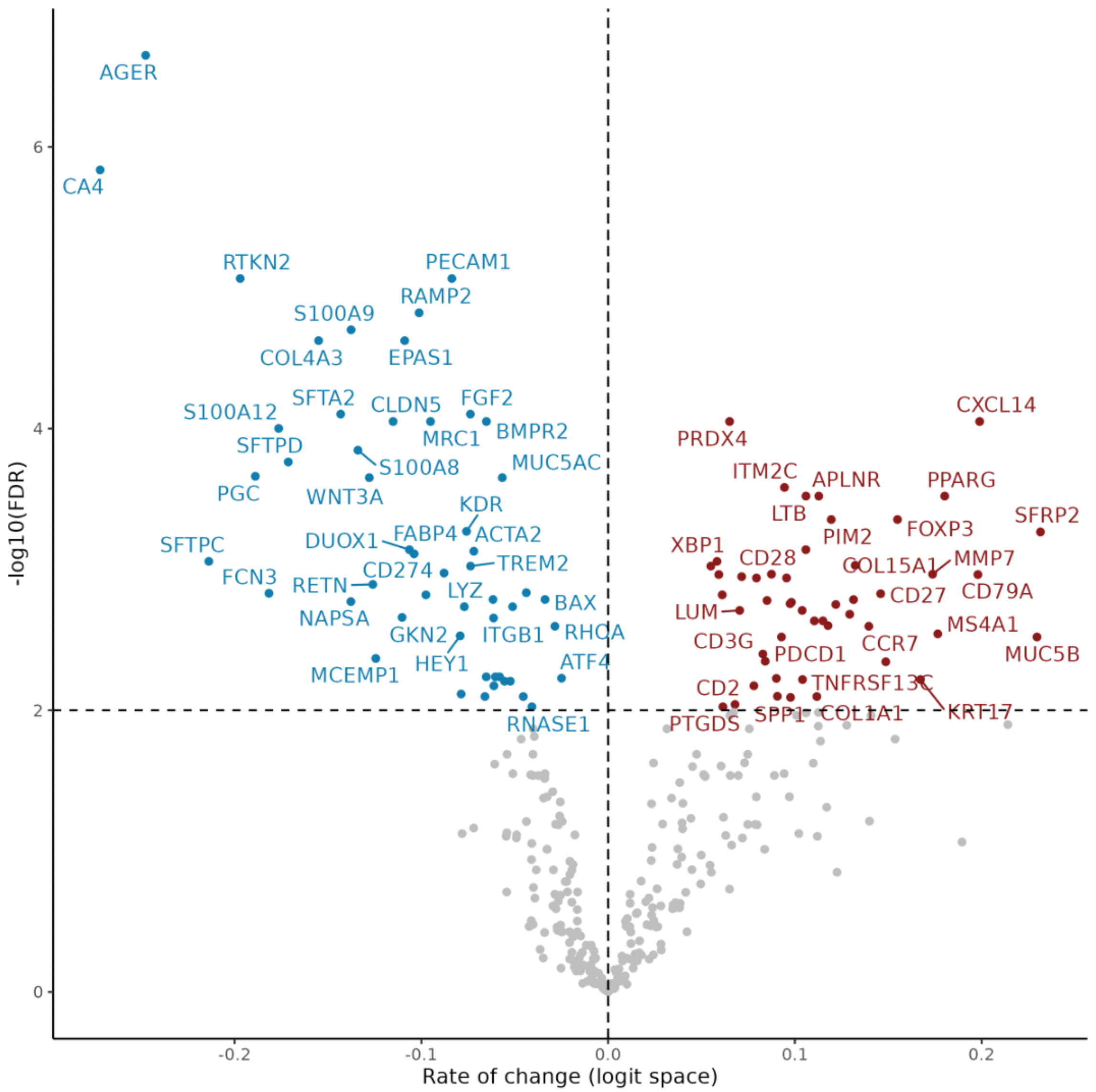


**Supplementary Figure 8: Gene expression changes associated with pathology score.** Gene expression was aggregated per sample and converted to proportions. Linear models were fitted to each gene after logit transformation of gene proportions against pathology scores. Significant genes (FDR < 0.01) are colored and labeled.

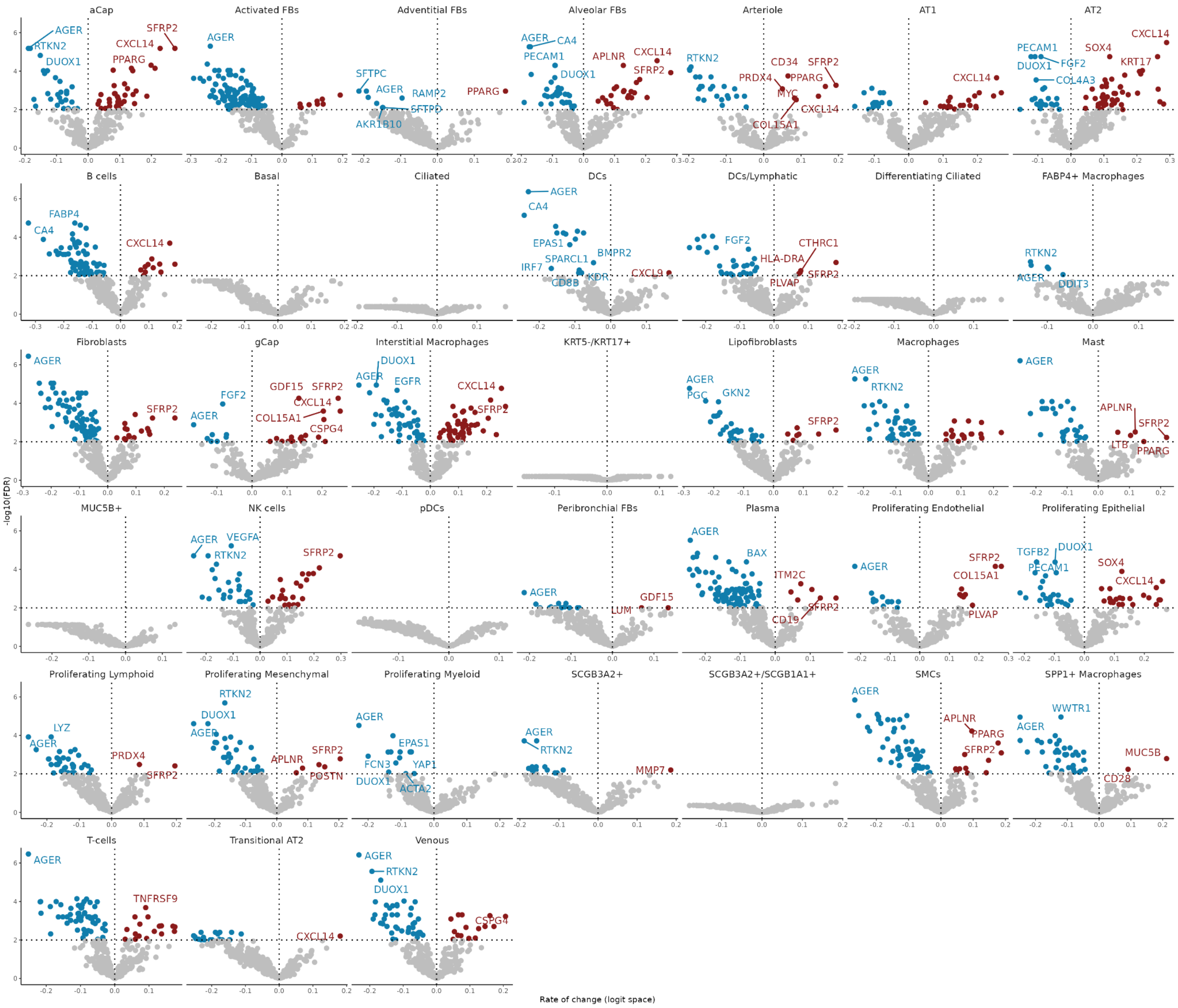


**Supplementary Figure 9: Cell type-level gene expression changes associated with pathology score.**

Complementary analysis to **Supplementary Fig. 8**, identifying gene expression changes associated with pathology score across each cell type. Significant genes (FDR < 0.01) are colored and labeled. Note: the limited precision of cell segmentation (leading to "contamination" of RNA from adjacent/overlying cells) contributes to some observed signals (for example, the AT1 lineage restricted gene *AGER* appearing as downregulated in numerous cell types). In this case, AT1 cell loss results in a reduction of contaminating gene expression in diseased samples. Conversely, increased *MUC5B* expression in *SPP1*+ macrophages reflects compositional differences in adjacent irregularly shaped airway epithelial cells.

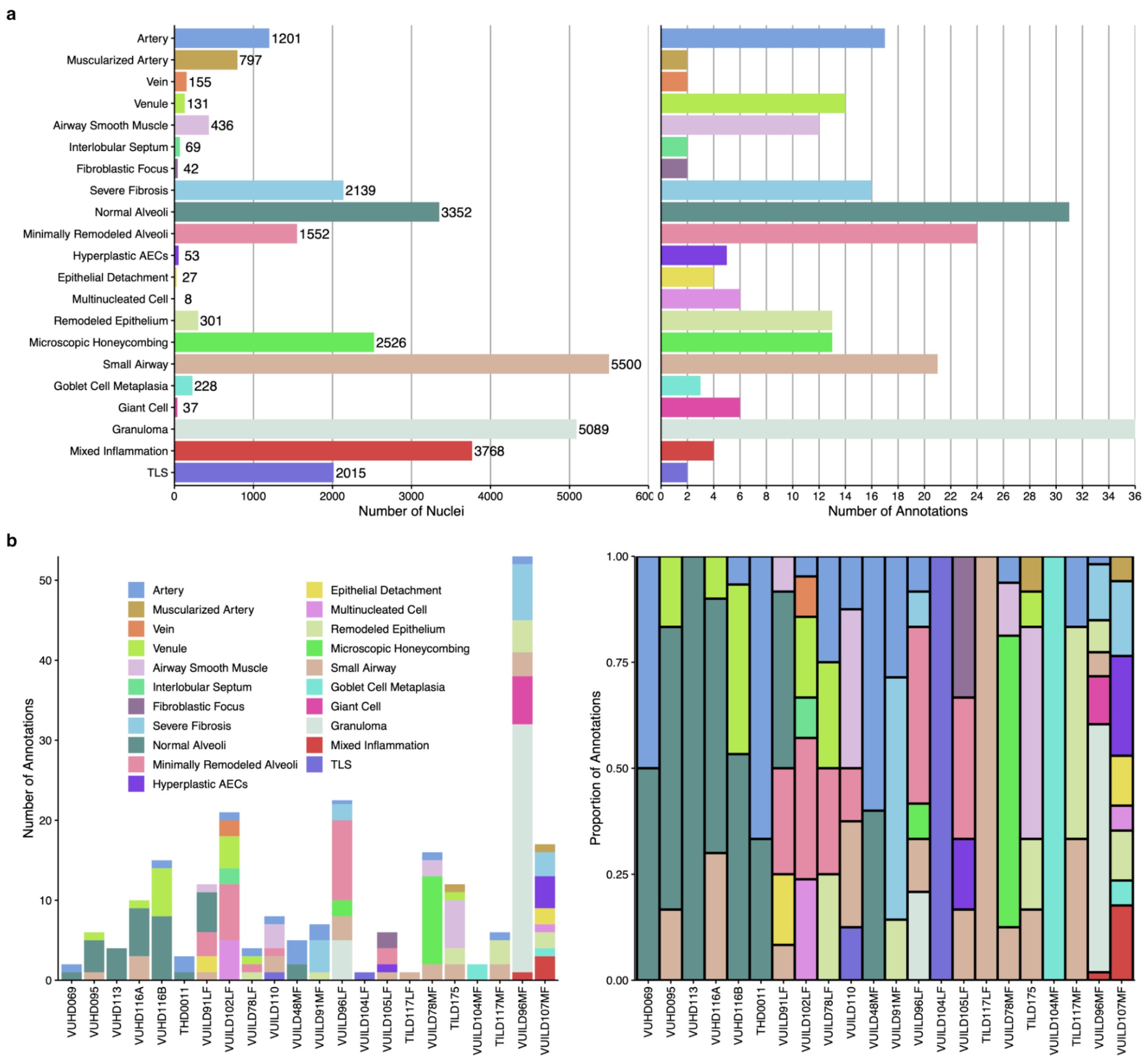


**Supplementary Figure 10: Distribution of annotations across samples.**
**a**, The number of nuclei per annotation (left) and total number of annotations of each type (right) across all samples. **b**, Distribution of annotation types across samples, both as counts (left) and as proportions (right) of the total number of annotations for a sample.

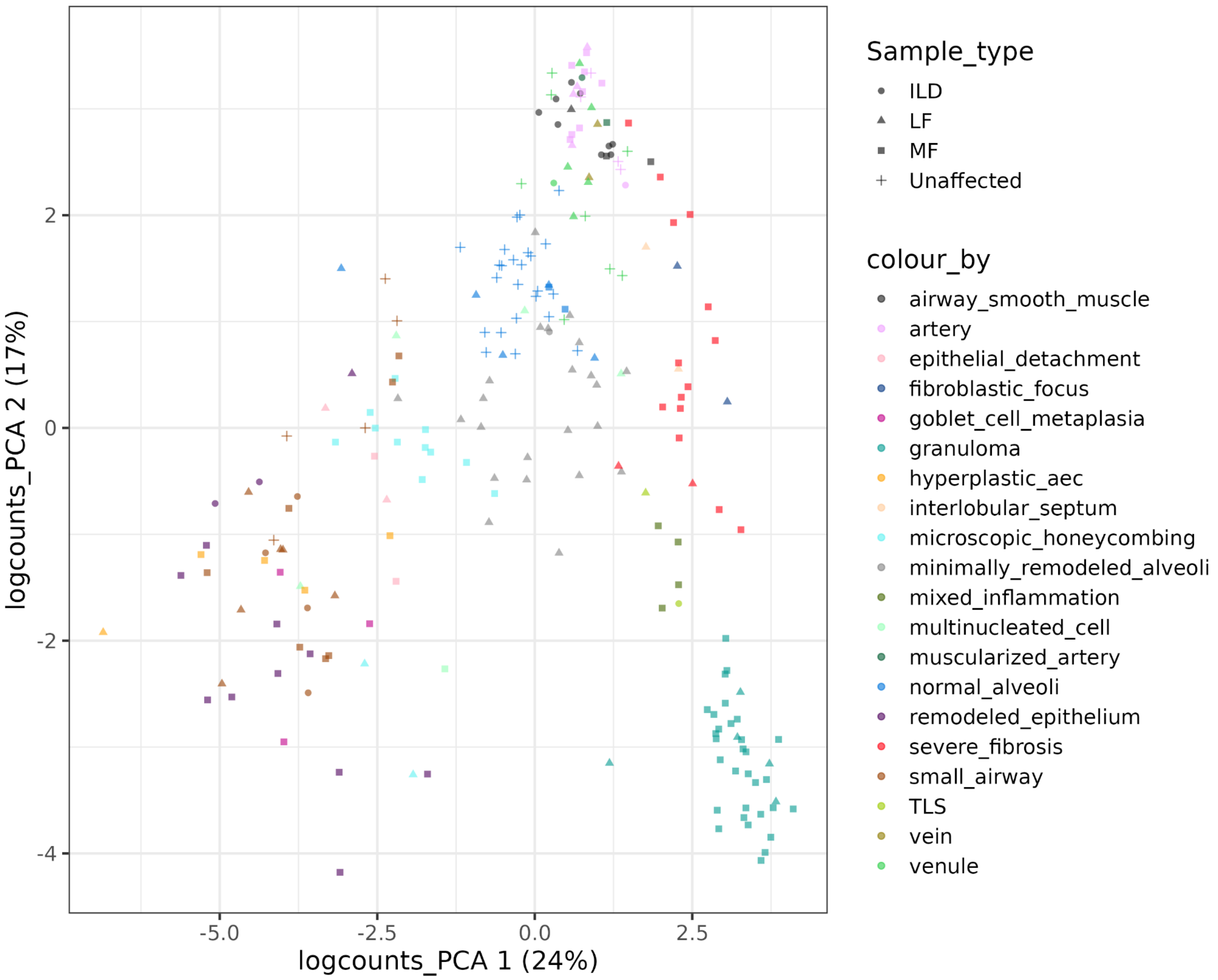


**Supplementary Figure 11a: PCA of gene expression patterns of annotated histological features.**

The gene expression profile of an annotation instance was constructed by aggregating the gene expression of cells assigned to the annotation. Gene expression per annotation instance was normalized using $log_2$ transformation after adding pseudo-count 1 and scaled by the size factor (the number of aggregated cells per annotation instance). PCA plot of histological annotations is coloured by annotation type using the $log_2$ normalized expression.

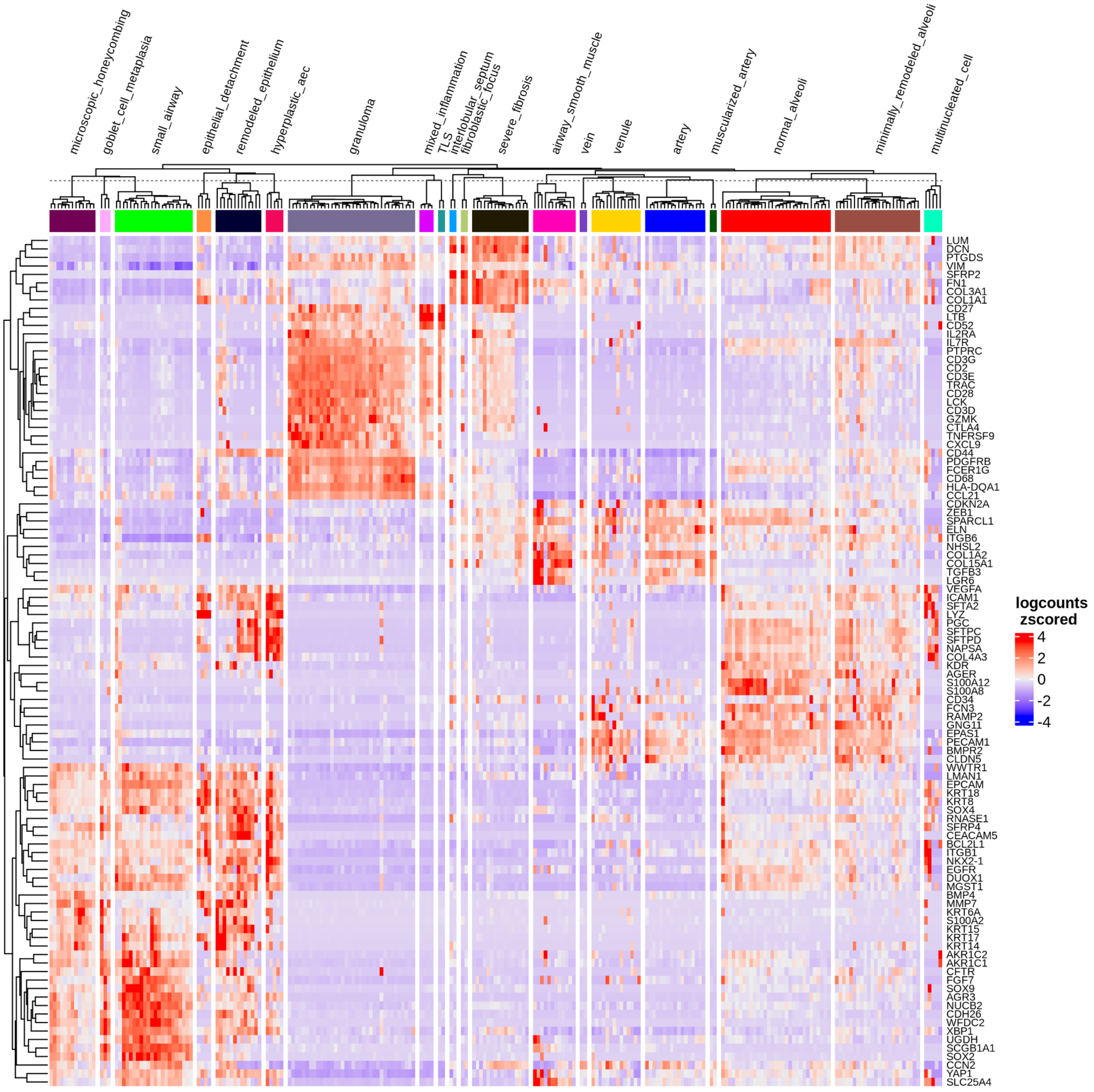


**Supplementary Figure 11b: Comparison of gene expression patterns underlying all annotated histological features.**

Differential gene expression was detected among all annotation types using generalized linear models implemented in *limma*[4] using the $\log_2$CPM transformed gene expression by comparing each annotation to the rest of the annotation groups. The top 100 genes ranked by F-statistics obtained using *topTable* were visualized. The giant cell annotation was removed for this analysis because the cells in this feature are included in the granuloma annotation.

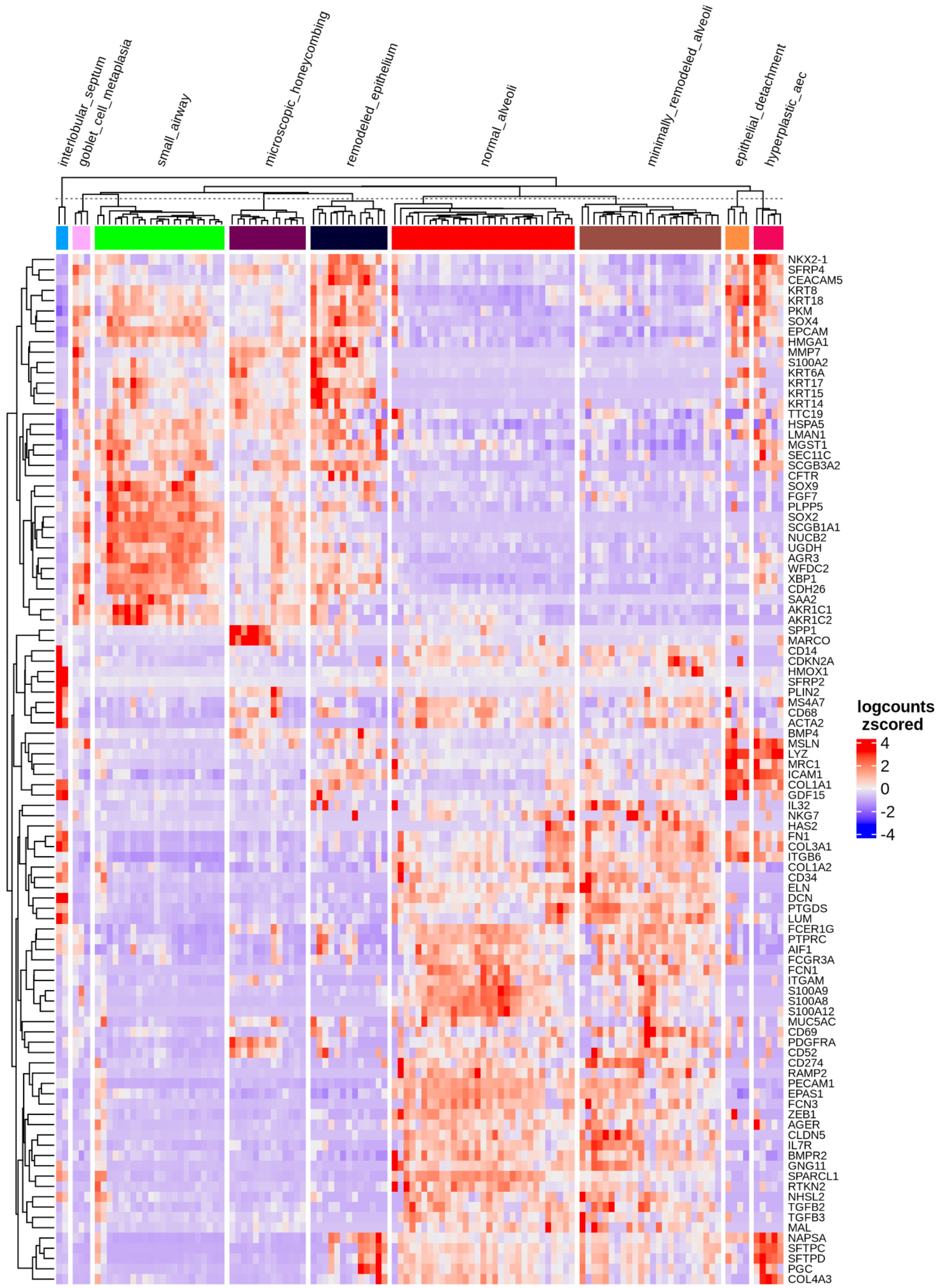


**Supplementary Figure 11c: Comparison of gene expression patterns underlying epithelial annotations only.**

Differential gene expression was detected among epithelial annotations using generalized linear models implemented in *limma*[4] using the $log_2$CPM transformed gene expression by comparing each annotation to the rest of the annotation groups. The top 100 genes ranked by F-statistics obtained using *topTable* were visualized.

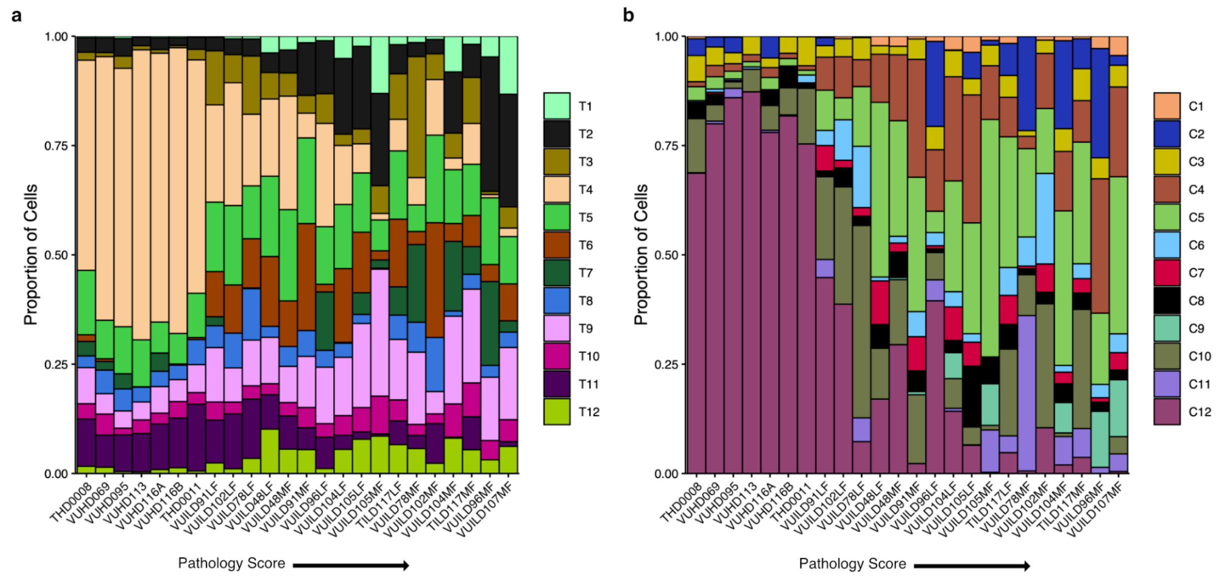


**Supplementary Figure 12: Niches assigned to each sample.** For each sample, transcripts were assigned to transcript-based niches using GraphSAGE[6] and cells were assigned to cell-based niches using Seurat v5[7]. Bar plots show the proportion of cells assigned to each transcript niche (**a**) and cell niche (**b**) per sample, ordered by pathology score from lowest to highest. For transcript niches (**a**), niches were determined for each cell by assigning cells to their closest hex bin after hex bin summarization as described in the methods.

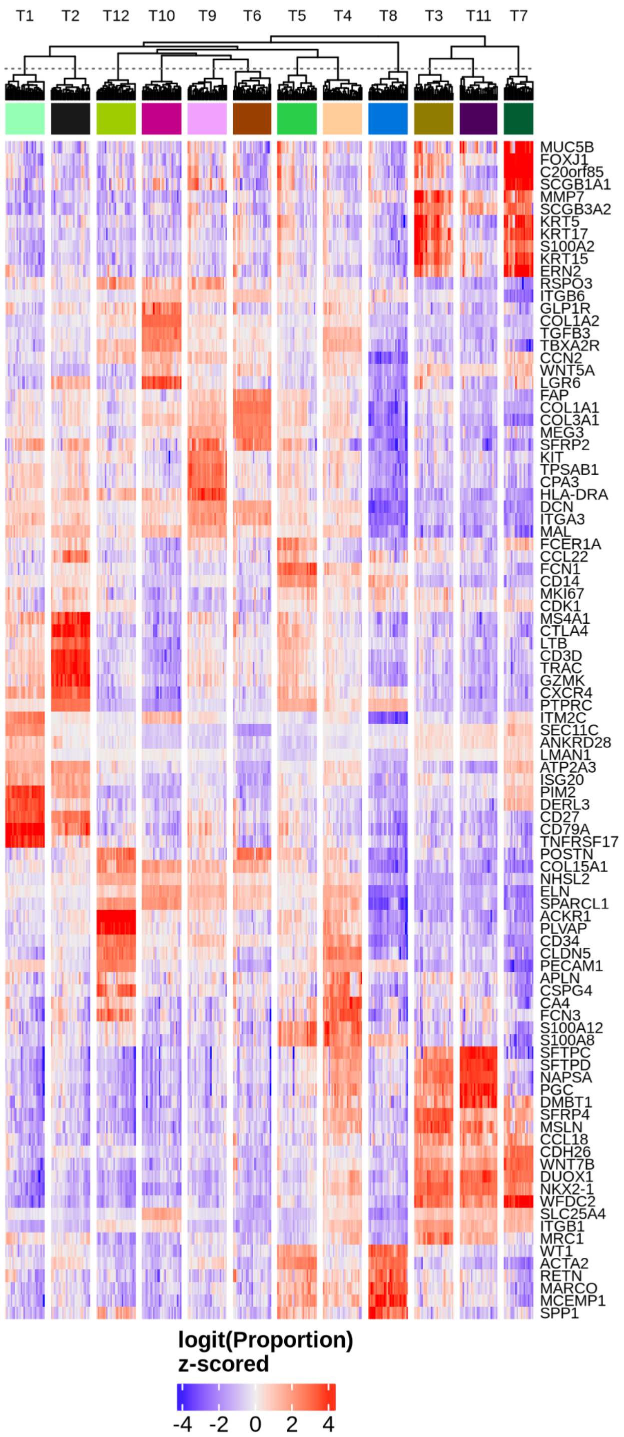


**Supplementary Figure 13: Transcript composition of transcript-based niches.** Differential proportion testing was performed for finding the representative genes per transcript niche using *propeller*[3] and *limma*[4]. For each transcript niche, gene proportions were calculated per sample, and contrasts were constructed to compare mean proportion in one transcript niche versus the other niches across samples. The top 5 up- and down-regulated genes were selected after ranking by proportion ratio and visualized in heatmaps.

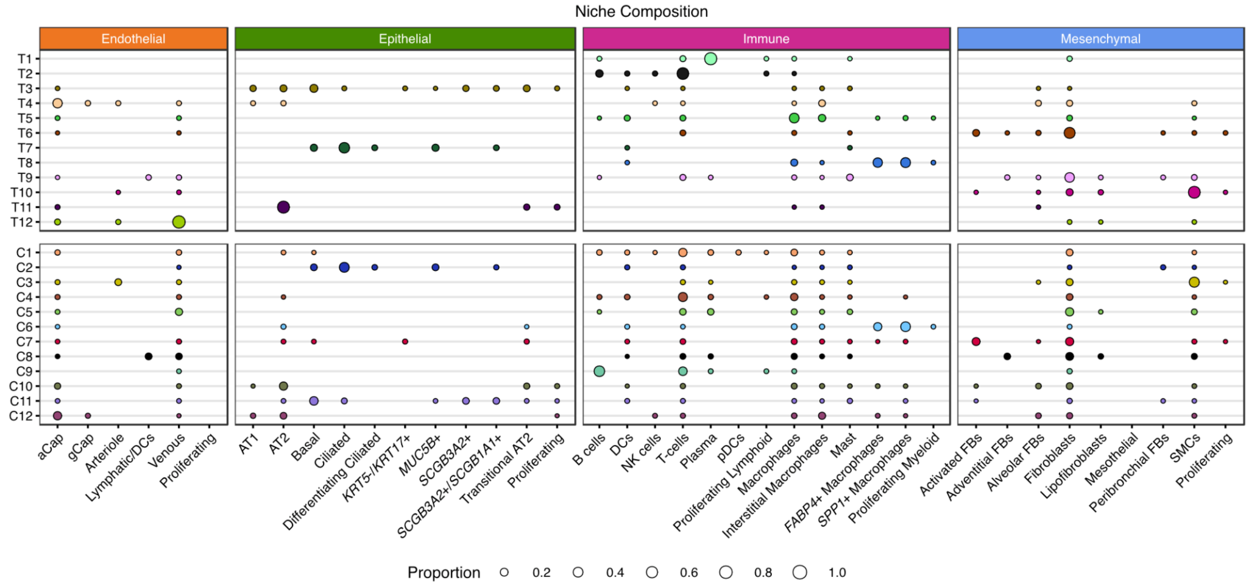


**Supplementary Figure 14: Cell type composition of niches.**

Cell type composition for transcript- (top) and cell-based niches (bottom), as a proportion of the total number of cells assigned to the niche (i.e., row sums). Proportions under 0.01 are not shown.

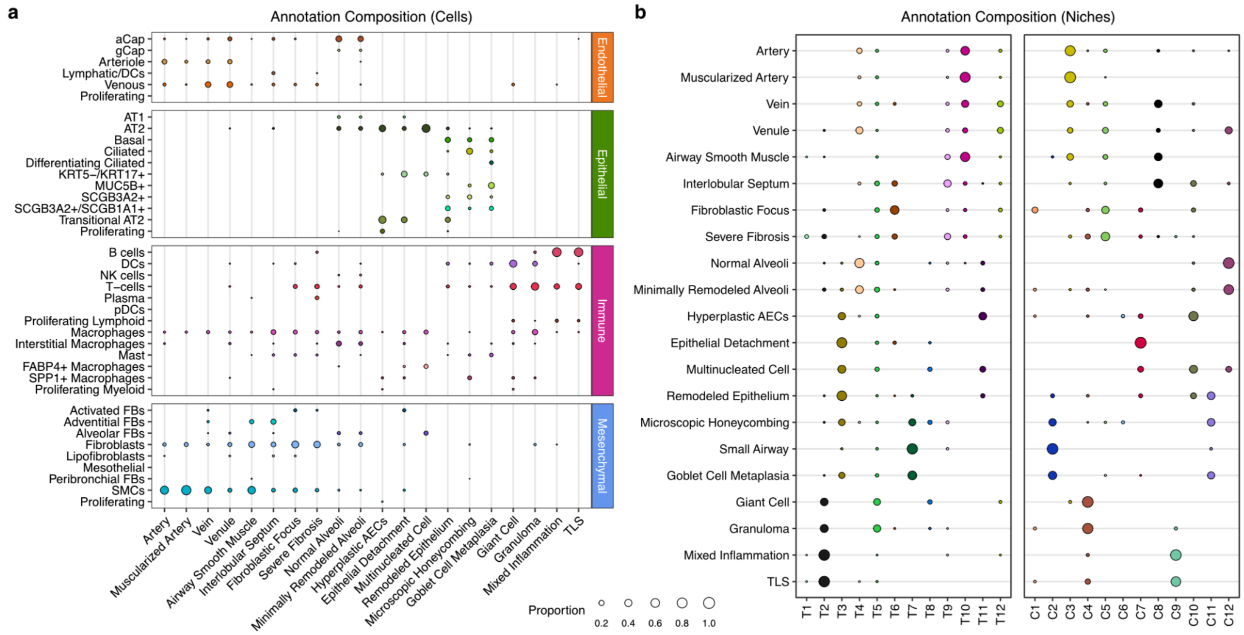


**Supplementary Figure 15: Cell types and niches in each annotation.**

**a**, Cell type composition of each transcript- (top) and cell-based niche (bottom), as a proportion of the number of cells assigned to that niche (each row sums to 1). This is a complete version of the plot shown in **Fig. 2c**. **b**, The niche composition of all annotations, as a proportion of the number of cells across an annotation (each row sums to 1). This is a complete version of the plot shown in **Fig. 3d**. For both **a** and **b,** proportions under 0.01 are not shown.

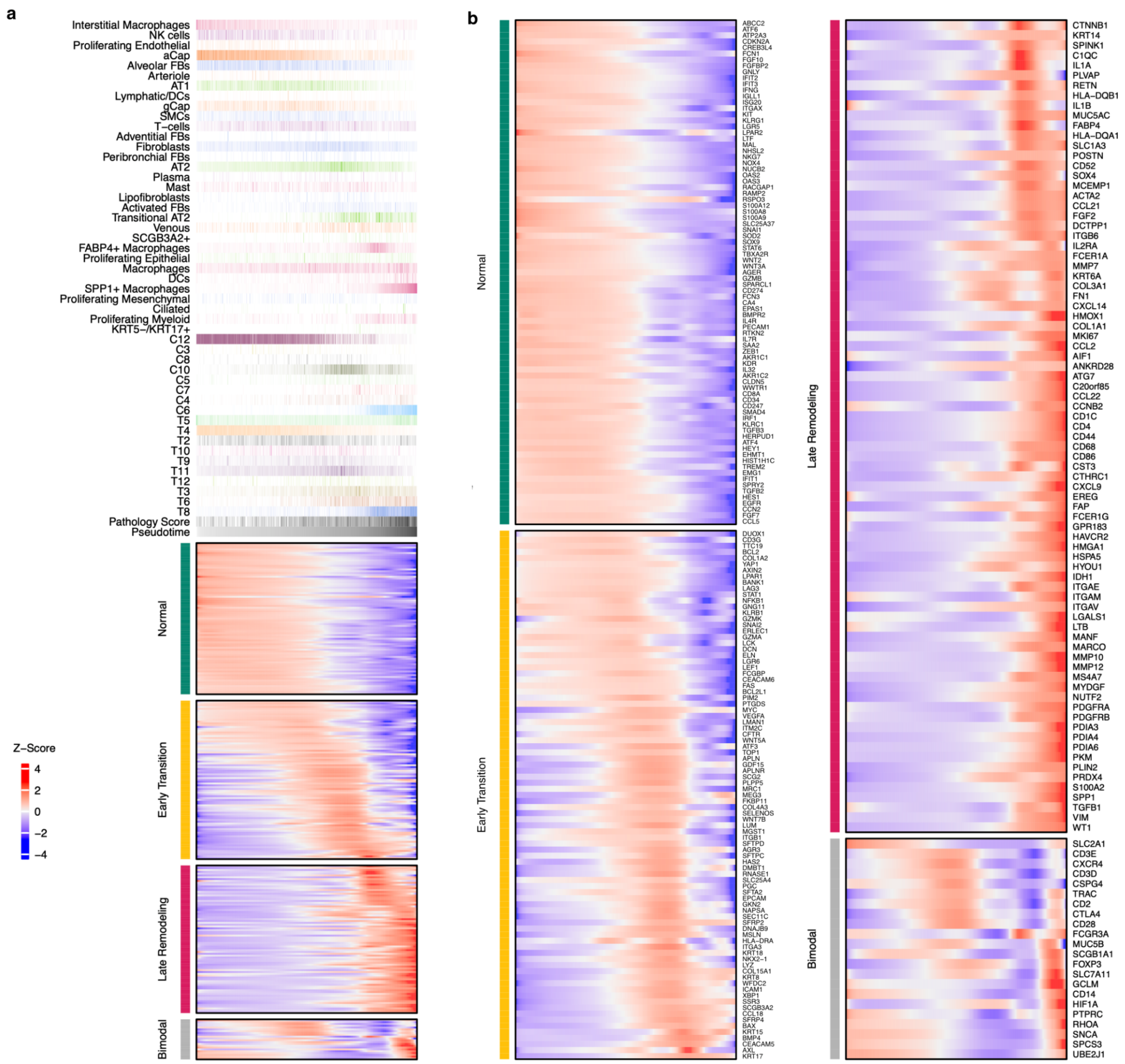


**Supplementary Figure 16: Heatmap of gene expression across airspaces along pseudotime.**

Expanded from **Fig. 6c**. **a**, In the heatmap, the predicted expression of each gene that was significantly associated with pseudotime, normalized and Z-scored, is depicted across the blue-red color gradient. Genes were divided into normal, early transition, and late remodeling based on their peak expression across the pseudotime trajectory. Genes were classified as bimodal if they peaked at multiple points along pseudotime, based on hierarchical clustering (see Methods). The top annotation shows all cell types, cell niches, and transcript niches that were associated with pseudotime (**Supplementary Fig. 17**, **18**), with the darkest shade of each color representing the maximum proportion of that cell type or niche found across all airspaces. Gene names for all categories are listed in **Supplementary Table 15**, and **b** shows the same heatmap at larger scale to aid in distinguishing individual genes on the plot.

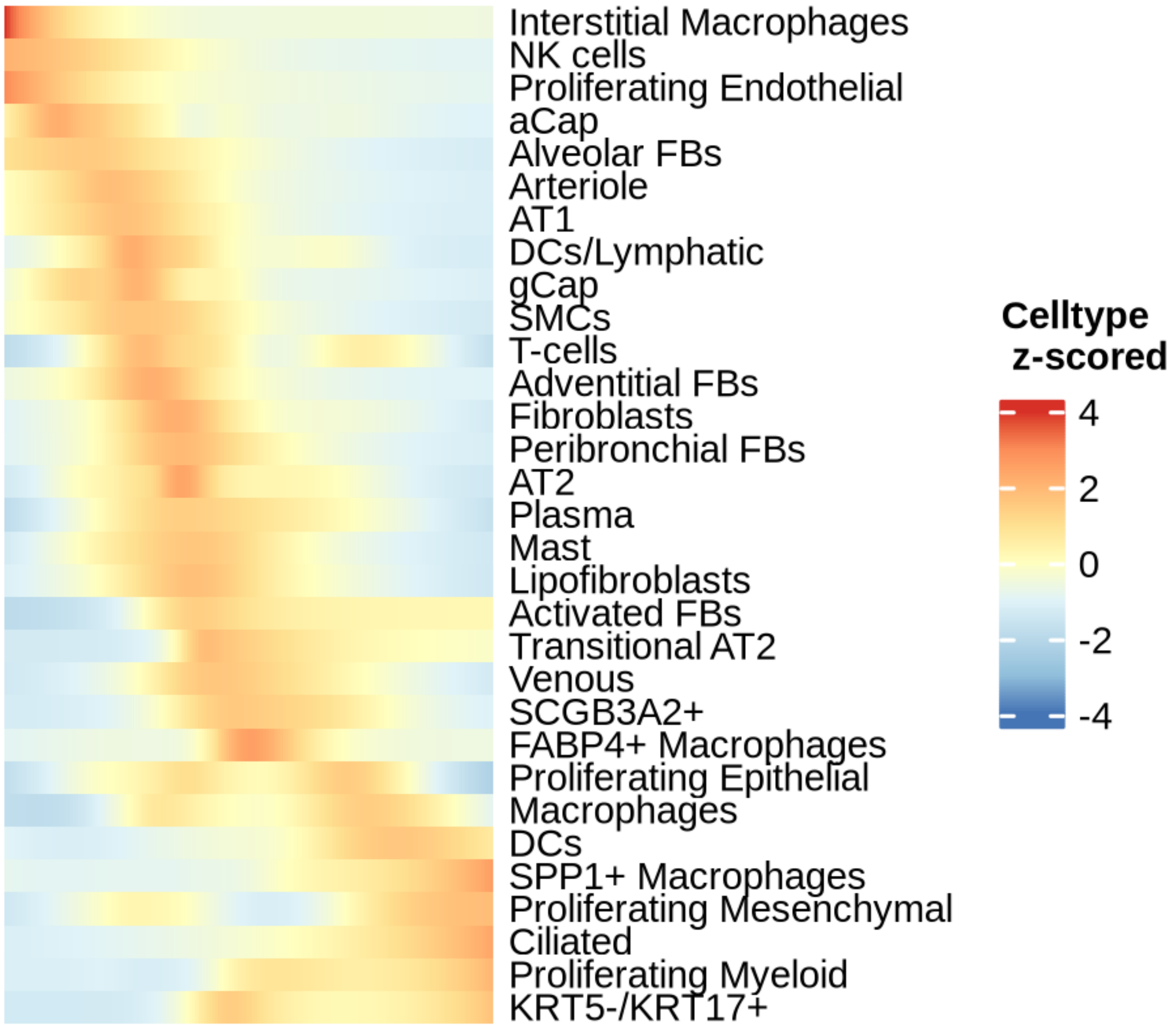


**Supplementary Figure 17: Cell type composition changes along pseudotime.**
Cell type proportion changes in alveolar air spaces along pseudotime. GAM was fitted per cell type with negative binomial distribution (knots=14) using cell type counts across lumens with log of total number of cells per lumen as offset. Cell types with > 0 count in 10 % of lumens were tested. Association of cell type abundance with pseudotime was tested using the *associationTest* function in *tradeSeq* with l2fc cutoff log2(1.5) and nPoints =10. Cell types with FDR < 0.01 were visualized in the heatmap.

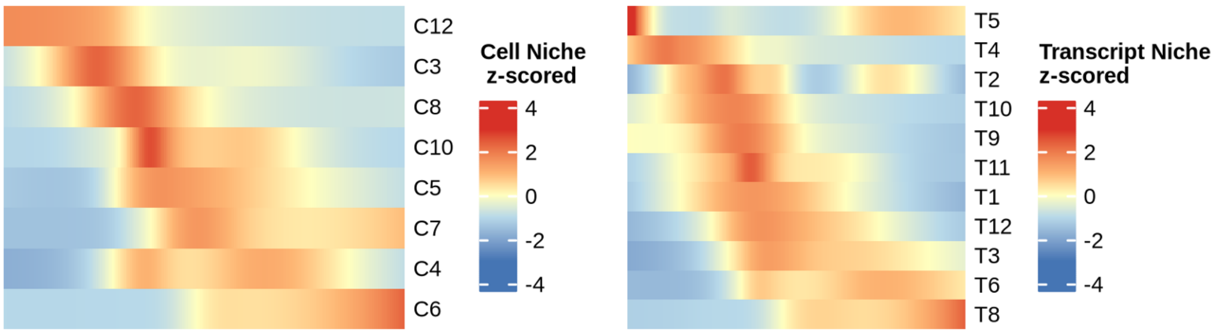


**Supplementary Figure 18: Cell and transcript niche composition changes along pseudotime.**

Cell niche proportion (**a**) and transcript niche proportion (**b**) changes in alveolar air spaces along pseudotime. GAM was fit per cell or transcript niche for niches that were found in at least 10% of alveolar spaces with negative binomial distribution (knots=9 for cell niches, knots=14 for transcript-based niches) using niche counts across lumens with log of total cell counts per lumen as offset. Association of niche abundance with pseudotime was tested using the *associationTest* function in *tradeSeq* with l2fc cutoff log2(1.5) and nPoints =10 for both niche types. Niches with FDR < 0.01 were visualized in the heatmap.

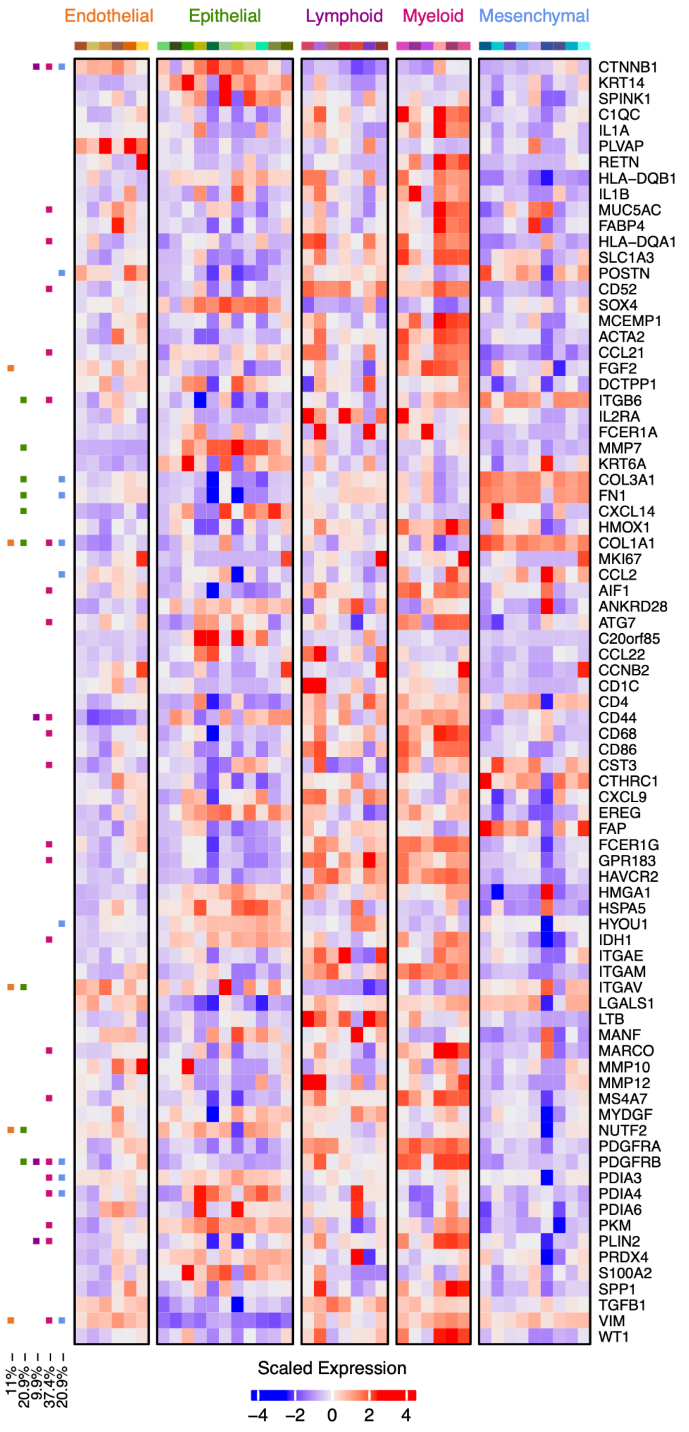


**Supplementary Figure 19: Contribution of cell types to changes in gene expression in late remodeling of alveolar airspaces.**

Heatmap shows scaled expression across the 81 genes with changes in gene expression in the late stage of alveolar remodeling as shown in **Fig. 6c**. Cell type colors (top) match those in **Fig. 3c** and **Supplementary Fig. 2**. On the left, boxes are filled in for each gene if it showed a significant change in expression in at least one cell type across the pseudotime of each of the following lineages: endothelial (orange), epithelial (green), lymphoid (purple), myeloid (pink), and mesenchymal (blue).

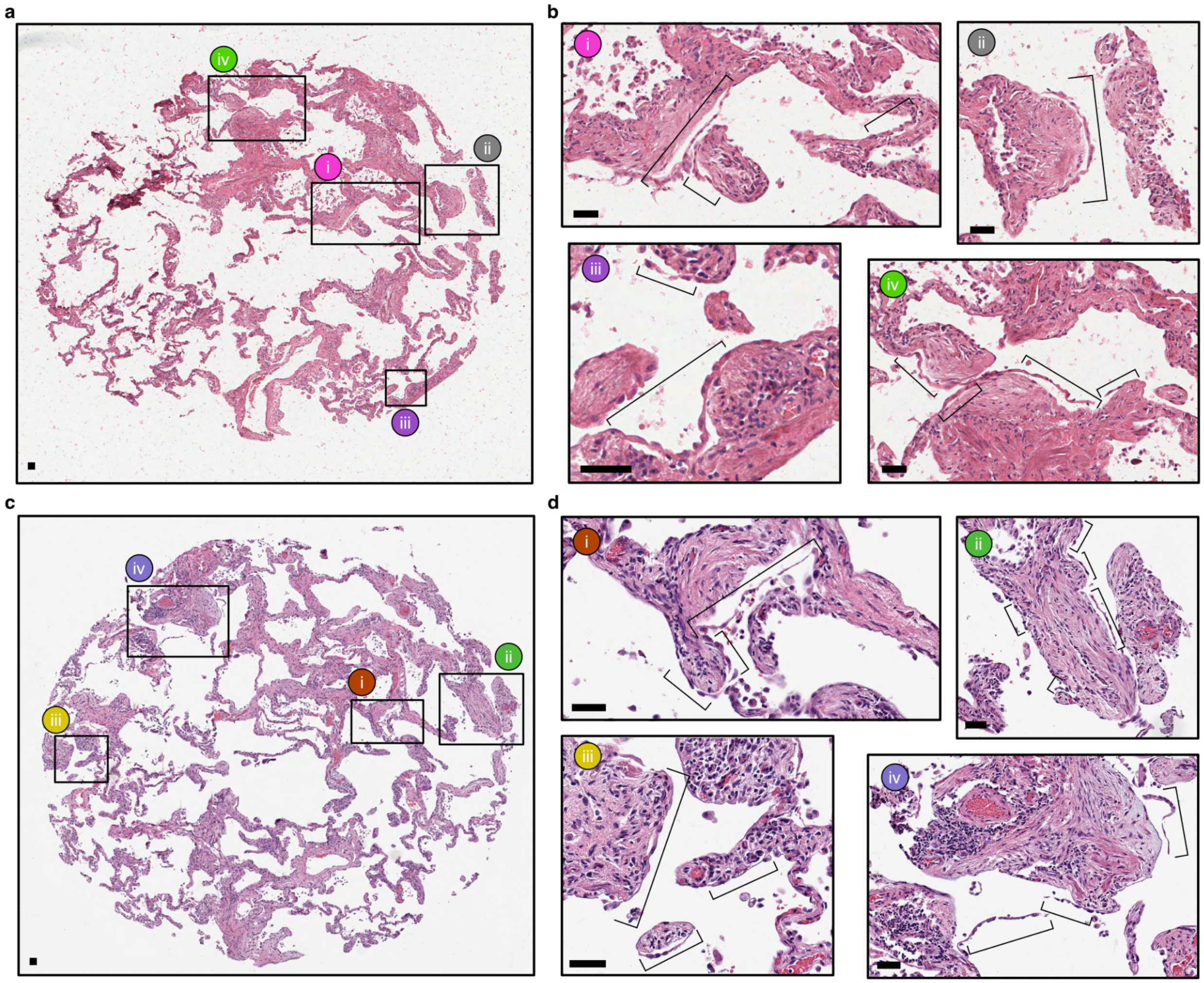


**Supplementary Figure 20: Epithelial detachment on a serial section that did not undergo Xenium processing.**

To verify that the epithelial detachment feature was not an artifact of Xenium processing, we took serial sections approximately 45 µm from the processed tissue section. H&E stains, shown here for sample VUILD48LF (NSIP diagnosis), show areas of epithelial detachment are present on both the original section (**a**,**b**) and the section did not undergo Xenium processing (**c**,**d**). **b** and **d** show the areas marked with boxes in **a** and **c**, respectively. Brackets denote epithelial detachment. Scale bars represent 50 µm.

1. Natri, H. M. *et al.* Cell type-specific and disease-associated eQTL in the human lung. *bioRxiv* (2023) doi:10.1101/2023.03.17.533161.
2. Sikkema, L. *et al.* An integrated cell atlas of the lung in health and disease. *Nat. Med.* **29**, 1563–1577 (2023).
3. Phipson, B. *et al.* propeller: testing for differences in cell type proportions in single cell data. *Bioinformatics* **38**, 4720–4726 (2022).
4. Ritchie, M. E. *et al.* limma powers differential expression analyses for RNA-sequencing and microarray studies. *Nucleic Acids Res.* **43**, e47 (2015).
5. Phipson, B., Lee, S., Majewski, I. J., Alexander, W. S. & Smyth, G. K. ROBUST HYPERPARAMETER ESTIMATION PROTECTS AGAINST HYPERVARIABLE GENES AND IMPROVES POWER TO DETECT DIFFERENTIAL EXPRESSION. *Ann. Appl. Stat.* **10**, 946–963 (2016).
6. Hamilton, W. L., Ying, R. & Leskovec, J. Inductive representation learning on large graphs. (2017) doi:10.48550/ARXIV.1706.02216.
7. Hao, Y. *et al.* Dictionary learning for integrative, multimodal, and scalable single-cell analysis. *bioRxiv* 2022.02.24.481684 (2022) doi:10.1101/2022.02.24.481684.
